# Genetic gradual reduction of OGT activity unveils the essential role of O-GlcNAc in the mouse embryo

**DOI:** 10.1101/2024.04.24.590926

**Authors:** Sara Formichetti, Agnieszka Sadowska, Michela Ascolani, Julia Hansen, Kerstin Ganter, Christophe Lancrin, Neil Humphreys, Mathieu Boulard

## Abstract

The reversible glycosylation of nuclear and cytoplasmic proteins (O-GlcNAcylation) is catalyzed by a single enzyme, namely O-GlcNAc transferase (OGT). The mammalian *Ogt* gene is X-linked and it is essential for embryonic development and for the viability of proliferating cells. We perturbed OGT’s function *in vivo* by creating a murine allelic series of four single amino acid substitutions reducing OGT’s catalytic activity to a range of degrees. The severity of the embryonic lethality was proportional to the degree of impairment of OGT’s catalysis, demonstrating that the O-GlcNAc modification itself is required for early development. We identified milder hypomorphic *Ogt* alleles that perturb O-GlcNAc homeostasis while being compatible with embryogenesis. The analysis of the transcriptomes of the mutant embryos at different stages suggested a sexually-dimorphic developmental delay caused by the decrease in O-GlcNAc. Furthermore, a mild reduction of OGT’s enzymatic activity was sufficient to loosen the silencing of endogenous retroviruses *in vivo*.

## Introduction

Thousands of nuclear and cytoplasmic proteins are reversibly modified at specific serine and threonine hydroxyls by the covalent linkage of the monosaccharide O-linked-β-N-acetylglucosamine (O-GlcNAc) (Wulff-Fuentes et al, 2021). O-GlcNAc is the main form of intracellular protein glycosylation in animals and an ever-increasing number of studies in cell lines have implicated this posttranslational modification in the modulation of essential various cellular functions, including the cell cycle (Capotosti et al, 2011a), translation (Li et al, 2019; Shu et al, 2021) and glycolysis (Yi et al, 2012; Nie et al, 2020). In spite of O-GlcNAc’s pleiotropy, the homeostasis of this modification is regulated by a single pair of highly conserved enzymes: O-GlcNAc transferase (OGT) (Kreppel et al, 1997; Lubas et al, 1997) and O-GlcNAc hydrolase (OGA) (Gao et al, 2001), both encoded by a single gene in the majority of animals (Martin et al, 2022). The mammalian *Ogt* gene is X-linked, and it is essential for early development (Shafi et al, 2000). Furthermore, the maternal inheritance of an *Ogt-*null allele results in preimplantation lethality, even in the presence of a functional paternal *Ogt* gene in heterozygous female embryos (O’Donnell et al, 2004). This implies that OGT or O-GlcNAc (or both) are necessary for oocyte maturation or for the cleavage stage embryo, which relies on the maternal payload of molecules. The requirement of a functional maternal *Ogt* allele poses a major hurdle to loss-of-function genetic studies. Another hindrance to functional studies in mammals is the fact that *Ogt* is essential for proliferation and survival of dividing cells (Li et al, 2023), implying that even conditional knock-out alleles eventually lead to cellular lethality that can confound molecular phenotypes.

All the above obfuscated the knowledge of the biology of O-GlcNAc during mammalian development, leaving salient questions unanswered. Firstly, it is not demonstrated whether the essential function of *Ogt* in the early embryo resides in its catalytic activity. To our knowledge, the only study pursuing a separation of OGT’s functions showed that in cell culture the O-GlcNAc transferase activity of *Ogt* is essential for cell proliferation, but that non-catalytic functions also contribute to the cellular phenotype resulting from OGT’s absence (Levine et al, 2021). Secondly, besides the numerous studies in adult tissues and cellular models, the molecular roles of OGT and O-GlcNAc in the early mammalian embryo *in vivo* have barely been explored. A growing body of evidence points towards O-GlcNAc acting as a mediator connecting external cues to the control of development and growth, both at the cellular and organismal level. O-GlcNAc’s donor substrate is UDP-GlcNAc (Haltiwanger et al, 1990), the end product of the metabolic pathway called hexosamine biosynthetic pathway, which utilizes 2-5% of intracellular glucose (Marshall et al, 1991) and is sensitive to nutrient levels (Marshall et al, 1991; Swamy et al, 2016). In adult mice, O-GlcNAc is involved in regulating energy conservation (Ruan et al, 2014) and eating behavior (Lagerlöf et al, 2016). In diabetic pregnancy, it partly mediates the negative effect of hyperglycemia on the mouse embryo (Kim et al, 2017). *Ogt* is a facultative escapee of X chromosome inactivation (XCI), found to variably escape random XCI *in vitro* in differentiating cells (Gendrel et al, 2014; Chen et al, 2016) and imprinted XCI *in vivo* during embryonic development (Cheng et al, 2019; Andergassen et al, 2021; Borensztein et al, 2017). Imprinted XCI (iXCI) is the selective inactivation of the paternal X that starts in the preimplantation embryo at the 4-cell stage and from the blastocyst onward is maintained only in the extraembryonic tissues (Mak et al, 2004; Okamoto et al, 2004). Notably, *Ogt*’s escape from XCI is particularly prominent in the postimplantation extraembryonic ectoderm (ExE), the extraembryonic tissue which gives rise to the placenta (Cheng et al, 2019). The placenta is the embryonic interface to the maternal environment, providing the embryo with gas, nutrients and hormones. One of the few genetic studies on mammalian *Ogt* found that the resulting double dose of OGT in mouse female placentae, remarkably conserved in humans (Howerton et al, 2013), is responsible for higher resistance of female mouse embryos to maternal stress compared to males (Howerton et al, 2013; Howerton & Bale, 2014). In humans, single amino acid substitutions of *OGT* are associated with X-linked intellectual disability (XLID; OMIM #300997), which is a neurodevelopmental syndrome with heterogeneous symptoms that, in the case of OGT-XLID patients, always include decreased intellectual ability (IQ<<70), low birth weight, short stature, drooling, compromised language skills and often anatomical brain and body anomalies (Pravata et al, 2020). OGT-XLID mostly affects males, as random XCI mitigates the effects of the mutations in females by leading to mosaicism of wild type and mutated alleles (Pravata et al, 2020). Being X-linked and a facultative escapee, *OGT* is likely involved in other sexually-dimorphic traits.

O-GlcNAc modifies numerous DNA-associated proteins, counting RNA polymerase II (Kelly et al, 1993), chromatin modifiers (Boulard et al, 2020) and various transcription factors, including the pluripotency master regulators OCT4 and SOX2 (Jang et al, 2012). Notably, biochemical pull-downs identified Ten-eleven translocation (TET) enzymes among the main OGT interactors (Deplus et al, 2013; Ito et al, 2013; Chen et al, 2013; Vella et al, 2013). TETs catalyze the hydroxylation of 5-methylcytosine (5mC) to 5-hydroxymethylcytosine (5hmC), which is the first step of 5mC re-conversion to unmodified cytosine (C) as 5hmC is not maintained upon cell division (Tahiliani et al, 2009). DNA methylation is particularly dynamic during preimplantation development, and plays an essential role in the repression of parasitic DNA sequences known as retrotransposons (Greenberg & Bourc’his, 2019). Recent evidence implicates OGT and local O-GlcNAcylation of chromatin factors in stable retrotransposon silencing: OGT conditional deletion in mouse embryonic stem cells (mESCs) causes a low-magnitude upregulation of many retrotransposon families (Sepulveda et al, 2024). Furthermore, targeted de-GlcNAcylation at the promoters of the most active type of murine endogenous retroviruses, namely intracisternal A-particle elements (IAPs), led to their full-blown reactivation (Boulard et al, 2020). It is therefore tempting to speculate a causal relationship between O-GlcNAc and developmental epigenetic silencing, a hypothesis which remains to be tested *in vivo*. In the fly, where the first O-GlcNAc genetic study was performed, *Ogt* is necessary to silence homeotic genes during postimplantation development (Gambetta et al, 2009), through the O-GlcNAcylation of Polycomb (Gambetta & Müller, 2014). Whether this functional connection between O-GlcNAc and developmental gene expression is conserved in the mammalian embryo has also not yet been investigated.

All the above urge the creation of genetic models that overcome the cellular and embryonic lethality caused by the loss of *Ogt*’s function. To our knowledge, the only reported murine allele causing impaired OGT’s activity and compatible with life is *Ogt*-C921Y, a mutation found in OGT-XLID patients that reduces both OGT and O-GlcNAc levels and it is useful to study XLID etiology since the resulting mice share features with the human patients. In this study, we created a series of mouse models whose OGT’s catalytic activity is impaired to a range of degrees, thanks to the substitution of key amino acids in the catalytic core. By analyzing the inheritance of the *Ogt*-hypomorphic alleles, we discovered that OGT catalysis, hence the O-GlcNAc modification, is an essential property of OGT *in vivo*. Then, we leveraged two of these alleles to profile the transcriptional changes resulting from reduced O-GlcNAcylation on i. preimplantation development, where we provide the first *in vivo* evidence that a decrease in O-GlcNAc is followed by the transcriptional upregulation of retrotransposons, and ii. postimplantation, where we found a sexually-dimorphic defect in placental differentiation upon lower O-GlcNAc levels.

To study the function of OGT at specific developmental stages without the confounding accumulation of defects during previous stages, we additionally created a mouse model bearing an *Ogt*-degron system for the induced fast removal of the OGT protein. To this aim, we genetically implemented the AID technology, which has been successfully applied to mammalian cells (Nishimura et al, 2009; Nora et al, 2017) and more recently to the mouse (Macdonald et al, 2022). Although the system was inefficient in preimplantation embryos grown *ex vivo*, it was effective in primary mouse embryonic fibroblasts (MEFs), where we characterized the immediate transcriptional changes and their evolution over time after acute OGT depletion.

## Results

### Gradual reduction of OGT’s catalytic activity proportionally impacts embryonic survival

We leveraged structural and biochemical knowledge on OGT (Martinez-Fleites et al, 2008) and used the CRISPR-Cas9 genome editing in the zygote to create a series of *Ogt* hypomorphic mouse mutants bearing single amino acid substitutions in the catalytic domain: Y851A, T931A, Q849N, H568A (Fig. 1A,B). These mutations hinder to various extents OGT’s binding to its donor substrate UDP-GlcNAc or the mechanism of catalysis. They were previously shown to reduce OGT’s enzymatic rate (V_MAX_) *in vitro* to respectively 70%, 20%, 5% and 0% (Fig. 1C) and OGT’s enzymatic activity (defined as V_MAX_/K_M_) to 24%, 18%, 2%, 0% of that of the wild type (WT) protein (Martinez-Fleites et al, 2008). To our knowledge, there is no report on the impact of these mutations on cellular O-GlcNAc levels, and thus the developmental phenotypes resulting from their murine alleles were unpredictable. Notably, we observed a clear correlation between the degree of impairment of OGT’s activity and the severity of the developmental phenotype, as measured by the success of gene targeting in the zygote and the rate of female germline transmission of the mutant allele (Fig. 1D, first two columns). Specifically, no CRISPR-edited founders were obtained at weaning for the mutation of the catalytic base (H568A), while none of the four founders obtained for the second most disruptive mutation (Q849N) transmitted the mutation to their progeny (Lazarus et al, 2011a). For *Ogt*-T931A (retaining ∼20% of WT OGT catalytic rate and activity), the female F_0_ founders transmitted the mutation to the F_1_ generation, albeit at a much lower rate than expected, and only heterozygous females were recovered at weaning (postnatal day 21; Fig. 1D, third column, and Fig. 2A; chi-squared test’s p-value = 1.94 e^-06^). This phenotype prompted us to study *Ogt^T931A^* mutants at the blastocyst stage. The least disruptive mutation, namely Y851A, was maternally transmitted to the F_1_ generation at mendelian ratio. Of note, *in vitro* OGT(Y851A) retains only ∼24% of WT V_MAX_/K_M_ (similar to the embryonic lethal T931A), but ∼70% of WT V_MAX_. The severity of embryonic lethality being proportional to the decrease in V_MAX_ rather than in the binding affinity suggests that *in vivo* the availability of the donor substrate is not the limiting parameter. Hemizygous *Ogt^Y851A/^*^Y^ and homozygous *Ogt^Y851A/Y851A^* animals were seemingly healthy, but displayed sub-lethality at weaning (Fig. 1D, third column, and Fig. 3A). We used this allele to address the function of *Ogt* after implantation, focusing on the placenta, which plays a vital role in supporting embryonic growth.

**Figure 1.**
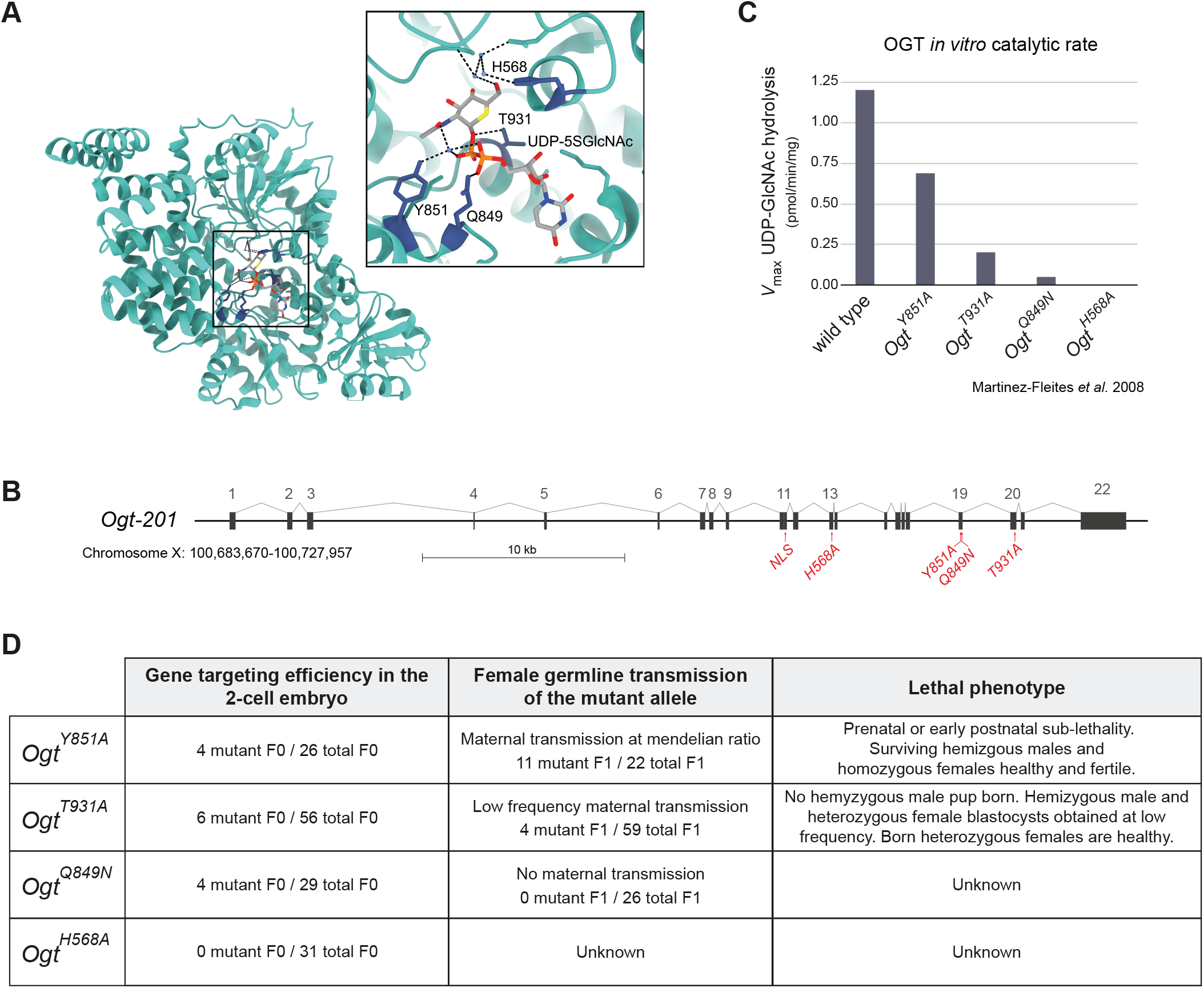
Structure-guided generation of *Ogt*’s hypomorphic allelic series. **(A)** Crystal structure of human OGT isoform 1 (UniProt O15294-1) in complex with UDP-5SGlcNAc (PDB 4GYY). The area in the square is shown as an insert with higher details. Except for two single amino acid substitutions outside the catalytic core, human OGT1 is identical to *Mus musculus* OGT isoform 1 (UniProt Q8CGY8-1), hence residues’ numbering is the same. The residues mutated in this study are shown in sticks representation in blue and UDP-5SGlcNAc in sticks representation coloured by heteroatom. T931 and Q849 establish direct interactions with the donor substrate. Y851 interacts with the donor substrate via hydrogen bonds with an intermediate water molecule. H568 is considered to be the catalytic base and in the crystal structure it coordinates different water molecules near the binding site. **(B)** Scheme of the murine *Ogt* genomic context with the location of the introduced point mutations. The exons of the longest transcript *Ogt-201* are numbered. **(C)** Catalytic rate measured *in vitro* for the four studied OGT single amino acid substitutions. Data are from Martinez-Fleites *et al*. 2008 (Martinez-Fleites et al, 2008) and for H568A also from Lazarus *et al*. 2011 (Lazarus et al, 2011b). **(D)** For the four hypomorphic mutants of OGT described in (**A**-**C**): number of animals (F_0_) bearing the correct point mutation after CRISPR-targeted mutagenesis in the zygote (founders); number of mutants (F_1_) produced by female founders (female germline transmission, i.e. maternal transmission); lethal phenotypes observed from the F_1_ generation onwards.

**Figure 2.**
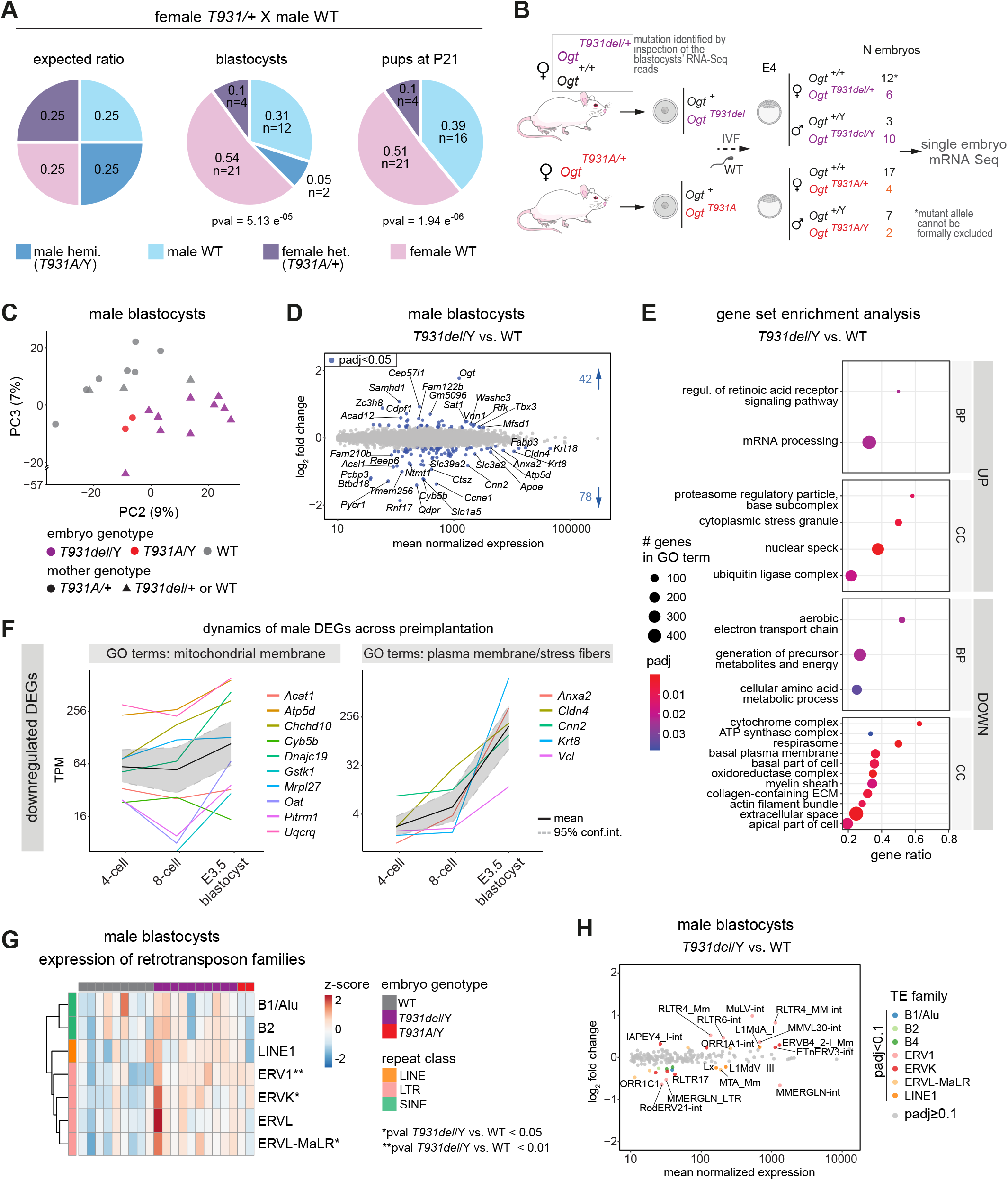
Maternal inheritance of a severe *Ogt* mutation causes preimplantation sub-lethality and perturbs retrotransposons silencing. **(A)** Number and frequency of genotypes for the progeny produced by the mating of female heterozygous *Ogt^T931A/+^* with WT males. The left pie chart shows the theoretical mendelian ratios for an X-linked mutation, the middle and right pie charts show the observed distribution of genotypes at the blastocyst stage and at weaning, respectively. P-values were computed using the chi-squared test for independence. **(B)** Experimental design for the study of blastocysts with maternal inheritance of a mutation at *Ogt*-T931. Oocytes from females *Ogt^T931A/+^* and seemingly *Ogt^+/+^* female littermates were fertilized *in vitro* with WT sperm and the embryos grown *ex vivo* to the blastocyst stage (E4), when they were collected for single embryo mRNA-Seq. For the embryos produced by *Ogt^T931A/+^* females, the sex and presence of the T931A mutation was determined by PCR genotyping of the cDNA and Sanger sequencing. For the rest of the embryos, the T931del allele was identified *a posteriori* by manual inspection of the RNA-Seq reads, and the embryos were genotyped based on sequencing reads mapping to *Ogt*. The WT genotype was assigned based on the absence of reads bearing a mutation at T931. The total number of embryos analyzed per genotype after the *in silico* filtering steps (Table S5) is indicated. Note that the absence of T931Adel-containing reads cannot formally exclude the presence of this allele within the group of heterozygous females. **(C)** Transcriptomes of individual male blastocysts produced by the experiment in (**B**) in the space defined by PC2 and PC3 of their principal component analysis (PCA), which result in a better separation based on embryonic genotype than PC1 and PC2 (shown in Fig. S2B). The variance explained by each PC is shown in parentheses. **(D)** MA-plot from DESeq2 differential expression analysis of single copy genes in *Ogt^T931A/^*^Y^ versus WT male blastocysts (N = 10 embryos per genotype). All genes with mean DESeq2-normalized gene counts > 10, adj. p-value < 0.05, any log_2_FC (DEGs) are colored, and their number is indicated. Genes standing out (and with abs(log2FC) ≥ 0.2) are labeled. **(E)** Gene set enrichment analysis (GSEA) of gene expression change in *Ogt^T931A/^*^Y^ versus WT male blastocysts. Among significant Biological Process (BP) and Cellular Component (CC) gene ontology (GO) terms, the first 20 based on Normalized Enrichment Score (NES) are shown. Terms are ordered based on gene ratio. The size of dots is proportional to the number of total genes of a GO term. Gene ratio = fraction of total genes of the GO term which are concordantly changing in mutant embryos. **(F)** Expression dynamics of *Ogt^T931A/^*^Y^ DEGs belonging to the indicated GO terms throughout preimplantation development (mRNA-Seq data from GSE66582 and GSE76505). The two biological replicates per stage were averaged. For each gene, the TPM value in the E3.5 blastocyst is the highest TPM value between the values in E3.5 ICM and E3.5 trophectoderm. The mean among all genes is drawn, as well as the 95% confidence interval, computed using basic nonparametric bootstrap (R function ‘mean.cl.boot’). Y-axis ticks are in log_2_ scale. TPM: Transcript Per Million. **(G)** Heatmap of the expression (in Fragments Per Kilobase Per Million, FPKM) of the main families of retrotransposons in the single male blastocysts produced in (**B**). Values are scaled by rows. Retrotransposon families are clustered based on Euclidean distance and coloured by class. P-values were computed using unpaired Wilcoxon rank sum exact test of FPKM values, and they are indicated when < 0.05. **(H)** MA-plot from DESeq2 differential expression analysis of retrotransposons in *Ogt^T931A/^*^Y^ versus WT male blastocysts (N = 9 WT, 10 *Ogt^T931A/^*^Y^ embryos). All repeats with mean DESeq2-normalized gene counts > 10, adj. p-value < 0.1, any log_2_FC are colored by their class. Repeats standing out are labeled.

**Figure 3.**
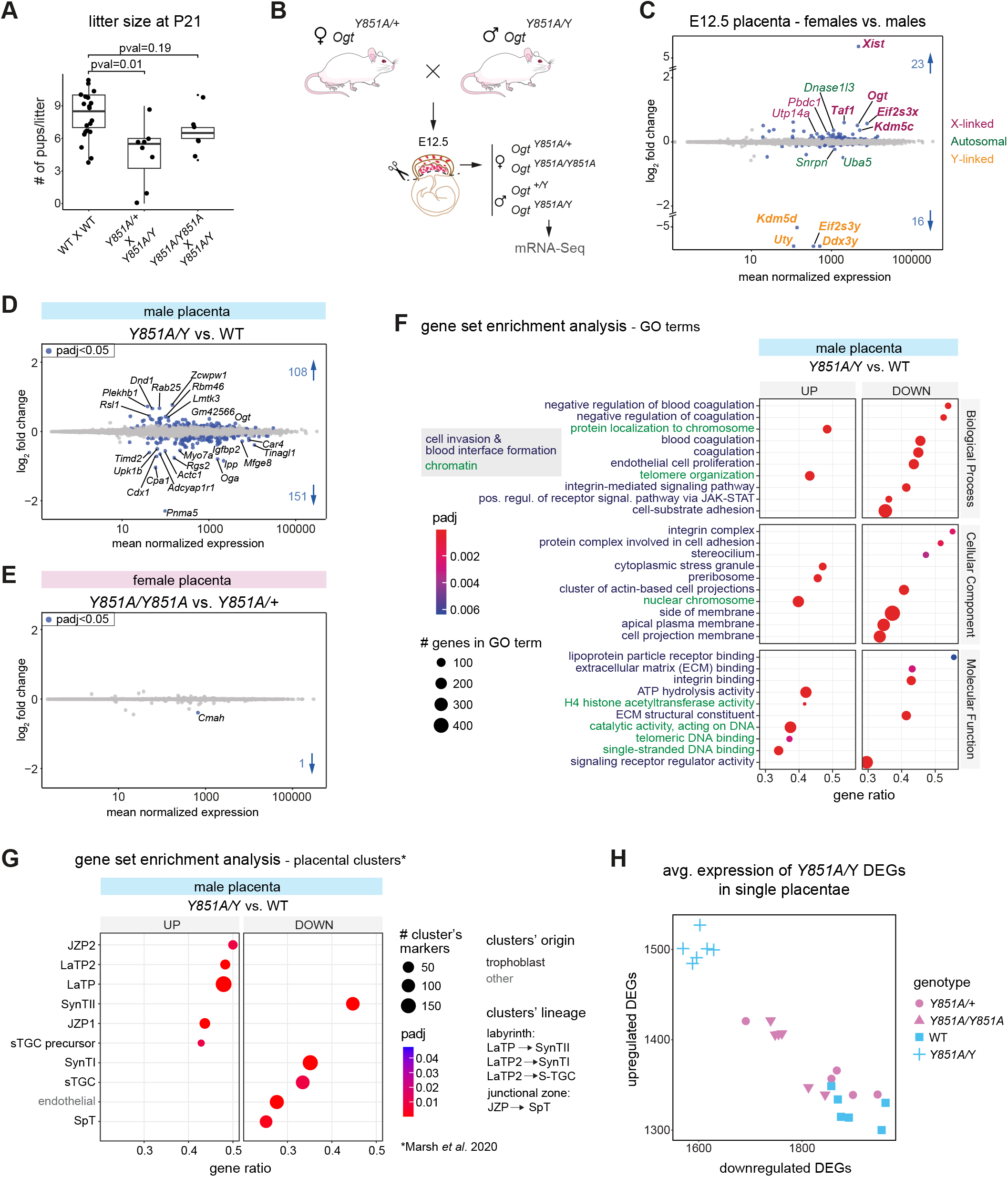
A mild reduction in OGT’s activity affects placental development in a sexually-dimorphic manner. **(A)** Litter size for intercrosses of WT mice and for crosses between mice bearing the *Ogt*-Y851A mutation. P-values are from unpaired two-sided Student’s t-test, assuming unequal variance. N = 20 WT crosses, 8 *Ogt^Y851A/+^* x *Ogt^Y851A/^*^Y^ and 6 *Ogt^Y851A/Y851A^* x *Ogt^Y851A/^*^Y^ crosses. **(B)** Breeding scheme used to produce placentae with the four different genotypes, which were analyzed via mRNA-Seq. **(C)** MA-plot from DESeq2 differential expression analysis of single copy genes in all female versus all male placentae in the dataset (all genotypes; N = 11 female, 12 male placentae). All genes with adj. p-value < 0.05, any log_2_FC are colored based on their location on sex or autosomal chromosomes, and their number is indicated. Genes standing out (and with abs(log_2_FC) ≥ 0.2) are labeled. Genes previously reported as sexually differentially expressed in E12.5-E18.5 placentae (Howerton et al, 2013) are in bold. (**D**,**E**) Separate DESeq2 analyses for female and male placentae, comparing (**D**) *Ogt^Y851A^***-**homozygous versus heterozygous female placentae or (**E**) hemizygous versus WT male ones. All genes with adj. p-value < 0.05, any log_2_FC are colored, and their number is indicated. Genes standing out (and with abs(log2FC) ≥ 0.2) are labeled. **(F)** GSEA of gene expression change in hemizygous *Ogt^Y851A/^*^Y^ versus WT male placentae. The first 10 GO terms for the three gene ontologies based on Normalized Enrichment Score (NES) are shown. Terms are ordered based on gene ratio. The size of dots is proportional to the number of total genes of a GO term. Gene ratio = fraction of total genes of the GO term which are concordantly changing in mutant embryos. JZP: junctional zone precursors; LaTP: labyrinth trophoblast progenitors; SynT-I: syncytiotrophoblast layer I; SynT-II: syncytiotrophoblast layer II; sTGC: sinusoidal trophoblast giant cells; SpT: spongiotrophoblasts. **(G)** Enrichment analysis of placental clusters’ marker genes (Marsh & Blelloch, 2020) expression changes in *Ogt^Y851A/^*^Y^ versus WT male placentae. All enriched clusters (adj. p-value < 0.05) are shown, ordered by gene ratio. The size of dots is proportional to the number of markers of each placental cluster. Gene ratio = fraction of total markers which are concordantly changing in mutant embryos. **(H)** Average DESeq2-normalized counts of DEGs, found in *Ogt^Y851A/^*^Y^ versus WT male placentae, in single placentae with all four genotypes analyzed. In (**D**-**H**), N per genotype = 6 placentae coming from at least two different litters, except female *Ogt^Y851A/+^* placentae for which one sample was excluded because outlier in unsupervised clustering.

Based on the sub-Mendelian inheritance of all catalytically hypomorphic alleles and on the severity of this phenotype being proportional to the reduction of OGT’s O-GlcNAcylation capacity, we conclude that the O-GlcNAc transferase activity of OGT, thus the O-GlcNAc modification itself, is essential for embryonic development, independently of the presence of the OGT protein.

In addition, we assessed *in vivo* the previously reported OGT’s nuclear localization signal (NLS) known as DFP_461-463_ (Seo et al, 2016a), by creating a murine allele where DFP_461-463_ is substituted by AAA (*Ogt^NLS-^*; Fig. S1A). This allele would have been useful to specifically investigate the functions of OGT in the nucleus. However, mutation of this putative NLS had no measurable effect on OGT nuclear localization, hence we found no evidence that the DFP motif is involved in OGT nuclear translocation (Fig. S1B,C).

### Maternal inheritance of a severe hypomorphic *Ogt* mutation causes preimplantation sub-lethality and perturbs retrotransposons silencing

*In vitro*, the T931A substitution causes an ∼80% reduction in OGT’s activity. *In vivo*, this mutation resulted in a significantly sub-mendelian distribution of the maternally inherited T931A mutated allele already at the blastocyst stage (Fig. 2A; chi-square test’s p-value = 5.13 e^-05^), where only 4 heterozygous females and 2 hemizygous males were found among 39 blastocysts from two F_0_ *Ogt^T931A/+^* females. Because of *Ogt* escaping imprinted (paternal) XCI, we would expect that female heterozygous can compensate for a hypofunctional maternal copy. Therefore, the preimplantation lethality of *Ogt^T931A/+^* heterozygous females, in other words the requirement of maternal inheritance of a catalytically active OGT, indicates that O-GlcNAc is essential either for oocyte maturation or in the cleavage stage embryo before the expression of the paternal allele.

The scarcity of blastocysts recovered bearing the T931A mutation prevented the statistical analysis of their molecular phenotype. However, because preimplantation development heavily relies on the maternal payload of RNA and proteins, we chose to assess the effect of the maternal genotype on the embryonic transcriptome. Thus, we performed single embryo mRNA-Seq (Picelli et al., 2014) on blastocysts generated through in vitro fertilization (IVF) of oocytes from a pair of F_0_ *Ogt^T931A/+^* females and from a pair of female littermates for which the *Ogt*-T931A mutation had not been detected by PCR; both groups were fertilized with the same WT sperm. Then, our manual inspection of the RNA-Seq reads mapping to *Ogt* unexpectedly uncovered the presence of an additional mutation of the T931 codon in the blastocyst population produced by the control females, namely a precise deletion of T931 (Fig. S2A). However, differently from the T931A substitution, the T931del allele was recovered in the blastocyst population with a high frequency (Fig. 2B), especially considering that *a posteriori* sequencing-based genotyping of the pair of control mothers revealed that only one of them was *Ogt^T931del/^*^+^ while the other WT. The data suggest that the deletion of threonine 931 is better structurally tolerated than its substitution by alanine, likely because of the next amino acid being also a threonine. This serendipity gave us the possibility to analyze the transcriptome of blastocysts with the precise deletion of an essential catalytic amino acid of OGT.

Principal component analysis revealed that the effect of the T931del mutation was visible only on the male transcriptomes (on PC3; Fig. 2C and Fig. S2B,C), most likely because at the blastocyst stage the heterozygous females express the WT paternal *Ogt*, which starts to be transcribed at early cleavage stages and then escapes imprinted X inactivation in the trophectoderm (Andergassen et al, 2021), as confirmed by higher levels of *Ogt* transcripts in WT female blastocysts compared to males (Fig. S2D). In good agreement with the PCA, a highly significant compensatory increase in *Ogt* expression is observed only in male hemizygous mutants and not in heterozygous mutant females (Fig. S2D). The upregulation of *Ogt* in response to decreased O-GlcNAc levels has been previously reported in cells and tissues (Qian et al, 2018; Willems et al, 2017; Pravata et al, 2019; Vaidyanathan et al, 2017; Park et al, 2017; Zhang et al, 2014).

We focused the analysis of gene expression changes on the males and found 120 differentially expressed genes (adj. p-value < 0.05, any log_2_FC, henceforth DEGs) in the *Ogt^T931del/^*^Y^ blastocysts when compared to the WT (from any maternal genotype), with 2/3 of the genes downregulated and 90% of the significant changes below 1 log_2_FC (Fig. 2D). Gene set enrichment analysis revealed the upregulation of proteasomal activity and stress granules along with the downregulation of amino acid metabolism, mitochondrial respiration and both transport and cell adhesion functions associated with the plasma membrane (Fig. 2E). This result is consistent with our recent finding that depleting nuclear O-GlcNAc from preimplantation nuclei slows down development, as the pathways downregulated in nuclear O-GlcNAc-depleted blastocysts included aerobic respiration and membrane transport (Formichetti et al, 2024). Thus, we tested whether DEGs due to the T931del mutation could be explained by a developmental delay by examining their dynamics across preimplantation development in unperturbed embryos. We reasoned that developmental gene expression changes should go in the opposite direction than gene misregulation if the latter is due to a delay. We found that the majority of the most significantly upregulated DEGs as well as many downregulated DEGs cannot be explained by a developmentally delayed transcriptome (Fig. S2E), while we cannot exclude this scenario for downregulated genes belonging to mitochondrial or cell adhesion-related GO terms (Fig. 2F). Of note, cell adhesion-related downregulated DEGs are normally highly expressed in the E3.5 trophectoderm, which again could be linked to a transcriptional delay of the mutant blastocysts or to a more pronounced effect of the lack of O-GlcNAc on this extraembryonic tissue (Zhang et al, 2017).

Previous works from us and others showed that the O-GlcNAc modification is required for stable silencing of retrotransposons in mESCs (Boulard et al, 2020; Sepulveda et al, 2024). However, it remains unclear whether this observation can be extended to other cell types or to the embryo and whether a milder and physiologically relevant perturbation of O-GlcNAc homeostasis could reactivate specific parasitic promoters. To assess the effect of reduced OGT’s activity on retrotransposon silencing during preimplantation development *in vivo*, we investigated their expression in the *Ogt^T931del/^*^Y^ blastocysts. Notably, most of the *Ogt^T931del/^*^Y^ embryos showed a low magnitude upregulation of retrotransposons belonging to families of long terminal repeats (LTRs) as well as long interspersed nuclear elements (LINEs) (Fig. 2G,H), while non-autonomous short interspersed nuclear elements (SINEs) contributed less to embryo clustering based on the mutation (Fig. S2F). Some single repeats showed downregulation, but they were generally lowly expressed and less robustly deregulated than upregulated repeats (Fig. 2H and Fig. S2G). Of note, the level of upregulation of retrotransposons in hypo-O-GlcNAcylated blastocysts is modest in comparison to that caused by inactivating mutations of *Trim28* (Boulard et al, 2020; Rowe et al, 2010) or dramatic DNA demethylation (Dahlet et al, 2020). Hence, our result could indicate that the residual level of O-GlcNAc contributes to retrotransposon silencing or that hypo-O-GlcNAcylation causes a global and incomplete disruption of heterochromatin rather than severely impacting the repressors of retrotransposons (e.g. HUSH and TRIM28 complexes or DNA methylation maintenance).

Because of hints of developmental delay of *Ogt^T931del/^*^Y^ blastocysts, we tested whether the deregulation of retrotransposons could be explained by their expression resembling that of an earlier stage’s transcriptome. The single repeats which are upregulated in mutant embryos are downregulated between the 8-cell and E3.5 stages in WT embryos, hence their altered level could be partly due to a developmental delay (Fig. S2H). However, the overall negative correlation between developmental expression changes of retrotransposons observed in WT embryos and changes due to the mutation is very weak, implying that a developmental delay is not sufficient to explain retrotransposon upregulation (Fig. S2H).

In summary, we found that the maternal inheritance of T931A is sub-lethal with high penetrance already during cleavage stages and characterized the transcriptome of blastocysts with a deletion of T931 which is compatible with preimplantation development. The male *Ogt^T931del/^*^Y^ transcriptomes revealed a translational stress response, the downregulation of metabolic and cell-adhesion functions, which can be partly explained by a potential subtle developmental delay, and a partial disruption of retrotransposon silencing.

### A mild reduction in OGT’s activity affects placental development in a sexually-dimorphic manner

The *Ogt*-Y851A substitution is compatible with life for hemizygous males and homozygous females, although sub-lethality was observed at weaning (Fig. 3A). This milder hypomorphic mutant enabled us to study the effect of a prolonged reduction in OGT activity on postimplantation development. We focused on the developing placenta, which was often found to be defective in embryonic lethal mouse mutants (Perez-Garcia et al, 2018). Moreover, *Ogt* escapes X-inactivation in the extraembryonic ectoderm (Andergassen et al, 2021; Cheng et al, 2019), resulting in the double dose of the enzyme in female placentae, which has been linked to higher sensitivity of male embryos to maternal stress (Howerton et al, 2013; Howerton & Bale, 2014) and could have other unexplored sexually-dimorphic developmental functions.

We crossed heterozygous *Ogt^Y851A/+^* females with hemizygous *Ogt^Y851A/^*^Y^ males and analyzed the transcriptome of single placentae dissected at E12.5 (Fig. 3B). In agreement with a previous study (Howerton et al, 2013), in WT and mutant samples a small fraction of genes was expressed in a sexually-dimorphic manner, whereby downregulated DEGs are mostly Y-linked and upregulated DEGs are mostly X-linked and include *Ogt* (Fig. 3C). We examined the impact of reduced OGT’s function on gene expression in the two sexes. We found a few hundred DEGs in hemizygous *Ogt^Y851A/^*^Y^ male placentae compared to their WT littermates (Fig. 3D) and only one in homozygous *Ogt^Y851A/Y851A^* females compared to heterozygous ones (Fig. 3E). Of note, DEGs found in hemizygous males were also dysregulated, although to a lesser non statistically-significant extent, in homozygous females; in addition, female heterozygous and WT males are indistinguishable based on the expression of DEGs (Fig. 3H). These observations are in agreement with a dose-dependent gradient of sensitivity to *Ogt* disruption.

The most significant downregulated gene sets in male *Ogt^Y851A/^*^Y^ placentae included *blood coagulation*, *endothelial cell proliferation* and many complexes and functions associated with the plasma membrane and the extracellular matrix (Fig. 3F). These gene sets point to the process of establishment of the maternal-fetal blood interface that is placental labyrinth (Woods et al, 2018). The E12.5 mouse placental labyrinth consists of two layers of multinucleated syncytiotrophoblasts (SynTI/II) that constitute the maternal-fetal gas and nutrient-exchange surface, and hormone-producing sinusoidal trophoblast giant cells (sTGCs); these three cell types differentiate from common labyrinth trophoblast progenitors (LaTPs) (Ueno et al, 2013). Between the labyrinth and the maternal decidua, the junctional zone hosts additional cell types with endocrine functions (Woods et al, 2018). To test the dysmorphism (or delay) in the formation of the different placental layers, we analyzed the expression of markers for all placental cell types spanning the E9.5 to E14.5 developmental window (Marsh & Blelloch, 2020). Notably, we found that markers for precursor cell types of both labyrinth and junctional zone (LaTPs and JZPs) were all upregulated in *Ogt^Y851A/^*^Y^ placentae, while markers for matured cell types (SynTs, sTGCs, SpT), which increase in percentage from E10.5 to E12.5 (Marsh & Blelloch, 2020), were all downregulated (Fig. 3G). Of note, downregulation was also observed for markers of endothelial cells, which also compose the fetal-maternal exchange barrier but are not of trophoblast origins (Fig. 3G). This transcriptional phenotype was highly penetrant among *Ogt^Y851A/^*^Y^ embryos, as shown by the expression of precursor and matured labyrinth markers in single placentae (Fig. S3A). Taking together the data above, we conclude that males with a reduced OGT activity display a delay in the differentiation of a functional embryonic placenta. This phenotype could be tissue-specific, for example due to *Ogt*’s role in the protein networks establishing cell-matrix contacts, or it could be associated with a slight developmental delay of the whole hemizygous *Ogt^Y851A/^*^Y^ embryo. We tested the latter possibility by staging *Ogt^Y851A/^*^Y^ embryos by morphometric analysis of limb development (Musy et al, 2018). This method has a precision of 0.5 day around E12.5 and did not reveal a difference between *Ogt^Y851A/^*^Y^ and WT embryos (Fig. S3B). However, we cannot exclude a delay of less than half a day.

Next, we assessed the impact of the Y851A mutation on the silencing of retrotransposons in the placenta. Hemizygous *Ogt^Y851A/^*^Y^ placentae but not homozygous *Ogt^Y851A/Y851A^* showed a very low magnitude upregulation of mRNAs transcribed from promoters of LTR and LINE classes of retrotransposons: LTRs of the endogenous retrovirus type K (ERVK) (*IAPLTR2_Mm*), two evolutionary recent LINE1 (*L1MdTf_I* and *L1MdTf_II*) and a full-length murine endogenous retrovirus-L (*MERVL-int*; Fig. S3C). These classes of retrotransposons are capable of retro-insertion (Gagnier et al, 2019) but we observed that the internal part (e.g. *Gag*, *Pol*) of IAPs, the most active autonomous LTR elements, was not significantly overexpressed in *Ogt^Y851A/^*^Y^ placentae (Fig. S3D). Thus, the epigenetic silencing mechanism that controls these elements is overall maintained upon a 30% reduction in OGT’s catalytic rate and *IAPLTR2_Mm* containing transcripts are likely produced by solo LTRs. Furthermore, we found a significant upregulation of two types of satellite repeats (IMPB_01 and MMSAT4), which are interspersed in the genome and very scarcely described in literature (Fig. S3E). It is noteworthy that male *Ogt^Y851A/Y^* placentae also upregulated gene sets related to chromatin remodeling and DNA binding (Fig. 3F). Heterochromatin being essential to control LINE1, LTRs and satellites (Montavon et al, 2021), a partial heterochromatin disruption could explain the increase in retrotransposons’ transcripts in the *Ogt*-mutant male placenta.

In conclusion, the data showed that a ∼30% reduction in OGT’s catalytic rate is sufficient to cause embryonic sub-lethality associated with male-specific delayed or defective placentation.

### Mouse alleles for fast inducible degradation of endogenous OGT

We also wanted to study the function of OGT at specific developmental stages without the confounding accumulation of effects during previous stages. To do so in the cleavage stage embryo, it is necessary to deplete the maternal payload of OGT molecules. Thus, we set out to achieve the acute removal of endogenous OGT protein *in vivo* in the mouse. We implemented a genetically encoded auxin-inducible degron (AID) system to the *Ogt* locus, by creating two knock-in alleles: i. the insertion of the *Oryza sativa TIR1* gene (*OsTIR1*) downstream of the ubiquitous promoter of the ROSA26 locus (Soriano, 1999) (ROSA26*^OsTIR^*allele, Fig. 4A) and ii. the N-terminal insertion of the minimal version of the AID peptide (44 amino acids) (Morawska & Ulrich, 2013; Macdonald et al, 2022) in frame with the longest isoform of *Ogt* (known as nucleocytoplasmic *Ogt* or *ncOgt*; *Ogt^NterAID-MYC-FLAG^* allele, henceforth *Ogt^AID^*, Fig. 4B).

**Figure 4.**
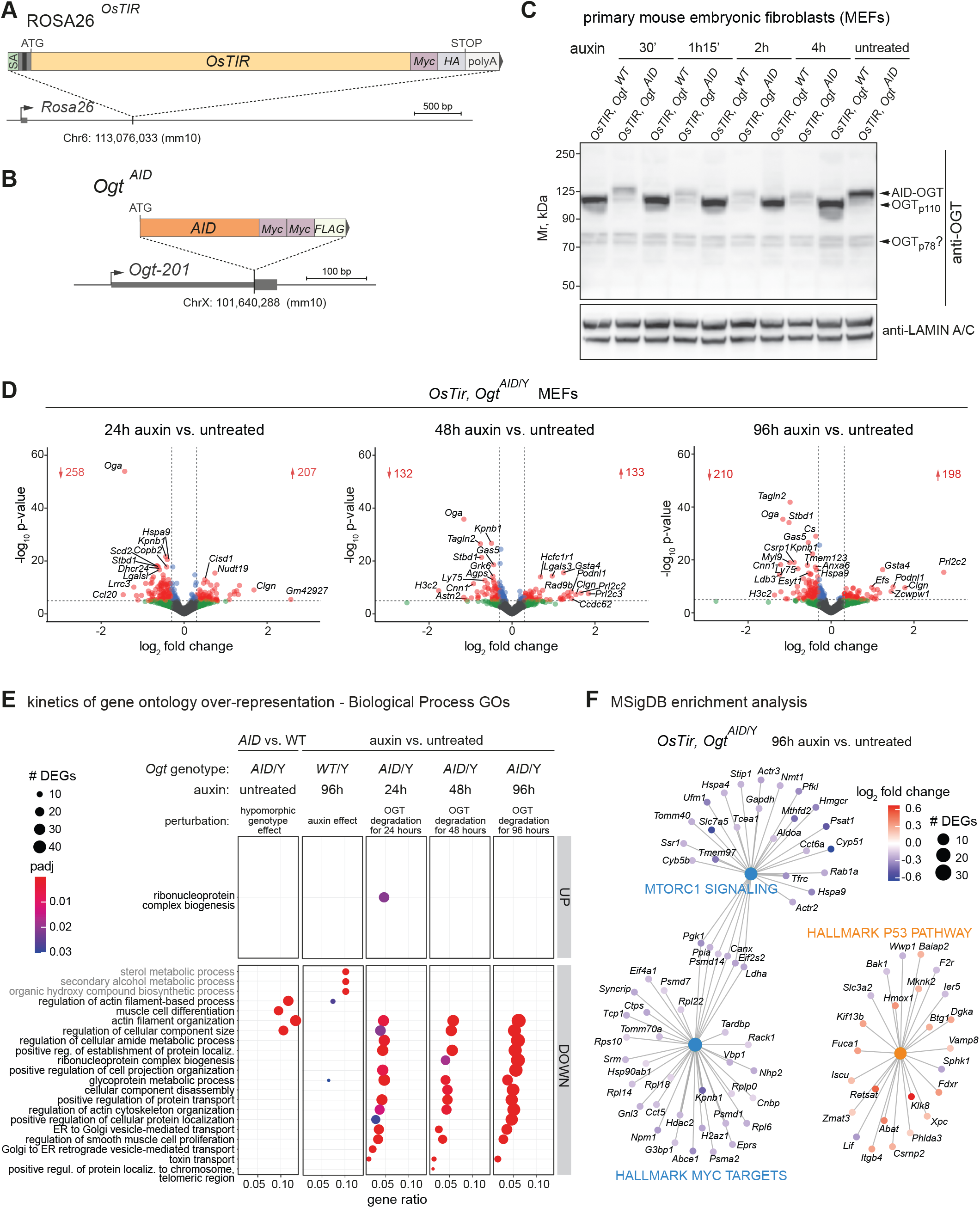
Kinetics of differential gene expression after rapid degradation of endogenous OGT in MEFs. **(A)** The ROSA26*^OsTIR^* allele was created by insertion of the coding sequence of *Oryza sativa TIR1* gene (*OsTIR1*) (fused to C-terminal Myc and HA adjacent tags) downstream the ubiquitous ROSA26 promoter. The transgene is integrated at the splicing acceptor (SA) site of ROSA26, resulting in ROSA26’s interruption and expression of *OsTIR1* from the endogenous promoter. **(B)** The *Ogt^AID^* allele consists of the targeted insertion of the minimal AID peptide’s sequence (44 amino acids) immediately downstream the initiating ATG codon of the longest *Ogt* isoform (*Ogt-201* or ENSMUST00000044475 or *ncOgt*), in frame with two Myc and one FLAG tag. Of note, anti-FLAG and anti-MYC antibodies did not give a specific signal for AID-OGT when tested using western blot or immunofluorescence, hence they were not used for any following experiment. **(C)** Western blot of endogenous OGT (ab177941 antibody) in whole cell protein extracts from male primary MEFs, untreated or treated with auxin for different amounts of time. From the same litter, an *OsTIR*,*Ogt^AID^*clone and a control *OsTIR*,*Ogt^WT^* clone were treated in parallel. Lamin A/C was probed as a loading control. **(D)** Volcano plots from DESeq2 analysis of gene expression changes in *OsTIR,Ogt^AID^* clones after 24, 48 and 96 hours of auxin treatment compared to untreated *OsTIR,Ogt^AID^* clones grown in parallel. Differentially expressed genes with p-value < 10^-5^ and absolute log_2_FC > 0.3 (i.e. 1.2-fold increase or decrease in expression) are colored in red and their number is indicated. **(E)** Gene ontology (GO) over-representation analysis of DEGs (adj. p-value < 0.05, any log_2_FC) from the same comparisons as in (**D**), plus untreated *OsTIR*,*Ogt^AID^* versus untreated *OsTIR,Ogt^WT^*clones (effect of the hypomorphic genotype), and *OsTIR,Ogt^WT^* control clones treated with auxin for 96 hours versus grown untreated (effect of the auxin drug). The first 25 most enriched Biological Process GO terms are shown, based on p-value across all comparisons. Terms are ordered by gene ratio. Gene ratio = genes belonging to the GO term / total number of deregulated DEGs for that comparison. UP = upregulated DEGs, DOWN = downregulated DEGs. Terms enriched due to auxin treatment on control clones are written in gray. **(F)** Analysis of enrichment for Hallmark gene sets of the Molecular Signatures Database (MSigDB) among DEGs after 96 hours of acute OGT depletion. For (**E**-**F**), DEGs after acute OGT depletion which were also differentially expressed in auxin-treated control cells were excluded before enrichment analyses of auxin-treated *Ogt^AID^*clones; DEGs with mean DESeq2-normalized counts < 10 across all samples were excluded from all enrichment analyses.

Heterozygous *Ogt^AID/WT^* females were viable at birth, healthy, fertile and transmitted the mutation at the expected ratio. However, hemizygous mutant males *Ogt^AID/^*^Y^ were never found at weaning, indicating that the addition of the AID tag to endogenous OGT impairs its stability or function *in vivo*. Nonetheless, we confirmed the expression of the *Ogt^AID/^*^Y^ allele in both male and female postimplantation mutant embryos (E7.5; Fig. S4A), therefore the *Ogt^AID^*allele is suitable to address OGT’s function at early embryonic stages.

We first tested the efficiency of the AID-OGT degron system in mouse embryonic fibroblasts (MEFs) derived from the breeding of females *Ogt^AID/WT^*with males homozygous for the *OsTIR* transgene (Fig. S4B). We analyzed the male *Ogt*-hemizygous clones, which only express either the WT or the AID-OGT protein. Of note, the presence of the AID causes a reduction of OGT protein amount of about half without auxin treatment (Fig. 4C and S4C), which could explain the lethality observed in hemizygous males. Upon the addition of auxin to the culture medium, the level of AID-OGT dropped to about 20% of that of WT after 30’ minutes (Fig. 4C and S4C). This reduction persisted for at least 4 days (Fig. S4D) and resulted in a global decrease of O-GlcNAc levels from 24 hours onwards (Fig. S4E).

### The AID-OGT degron system is inefficient in *ex vivo* grown preimplantation embryos

Having validated the OGT-degron system in MEFs, we applied it to our model of interest, namely the preimplantation embryo. We obtained zygotes with all relevant genotypes (i.e. females *OsTIR*,*Ogt^WT/WT^* and *OsTIR*,*Ogt^AID/WT^*; males *OsTIR*,*Ogt^WT/^*^Y^ and *OsTIR*,*Ogt^AID/^*^Y^) by IVF of oocytes derived from *OsTIR*-homozygous,*Ogt^AID/WT^* females or control *OsTIR*-homozygous,*Ogt^WT/WT^*littermates (always using WT sperm). Half of the embryos were cultured in presence of auxin from fertilization to the morula stage (embryonic day 2.5; Fig. S5A, top row). We had previously determined that the experimental concentration of auxin was not toxic to the embryos (Fig. S5B) and that the *OsTIR* transgene was expressed in blastocysts from *OsTIR*,*Ogt^AID/WT^* oocytes (Fig. S5C). Moreover, the rate of formation of expanded blastocysts from zygotes grown in auxin was not significantly different between *OsTIR*,*Ogt^AID/WT^*and control oocytes (Fig. S5D), hence all four possible genotypes should be equally represented in the population of morulae. OGT depletion is predicted to cause a reduction of global levels of O-GlcNAc in *OsTIR*,*Ogt^AID/^*^Y^ males (∼25% of the embryos), which do not bear any WT *Ogt* gene. We quantified the O-GlcNAc immunofluorescence (IF) signal in single morulae from *OsTIR*,*Ogt^AID/WT^*mothers after auxin treatment and found that the signal was only slightly diminished (Fig. S5E,F). Using the same breeding scheme, we also stained OGT in zygotes (3.5 hours post-IVF) cultured in auxin from fertilization. We chose this stage because the OGT signal is enriched in the paternal pronucleus (Fig. S5G, no auxin), enabling an easier qualitative assessment of OGT reduction. None of the auxin-treated zygote displayed a clearly diminished OGT signal, even with a higher concentration of auxin (Fig. S5G, N = 29).

However, we could not exclude a low level of auxin-induced OGT degradation below immunofluorescence sensitivity. Thus, we investigated whether a molecular difference emerged in auxin-treated *OsTIR*,*Ogt^AID/^*^Y^ embryos by means of transcriptomics. We collected zygotes recovered from crosses of *Ogt^AID/WT^* females with *OsTIR*-homozygous males, cultured them *ex vivo* for 72 hours, then treated half of the morulae with auxin for 24 hours and collected the blastocysts for single embryo mRNA-Seq (Fig. S5A, bottom row). Principal component analysis (PCA) showed a partial separation between the *Ogt^WT^*and *Ogt^AID^* genotypes only for males and only in the presence of auxin (Fig. S5H). Accordingly, gene expression changes were more pronounced for hemizygous males treated with auxin (Fig. S5I). The male-specific gene expression change induced by auxin treatment provides an indirect indication that the degron system is activated and partially impacts OGT’s function. It is noteworthy that, among the 32 upregulated DEGs, 4 were also significantly upregulated at a higher degree in *Ogt^T931del/^*^Y^ blastocysts: *Washc3,* which promotes actin polymerization at the surface of endosomes, the lysosomal transporter *Mfsd1*, mitochondrial insertase *Mtch2* and isocitrate dehydrogenase 1 (*Idh1*) (in bold in Fig. S5I). This further supports auxin-induced depletion of OGT, although to a mild extent. However, the resulting transcriptional change is of low magnitude and high variance, which results in a low number of DEGs (Fig. S5I) and prevents drawing conclusions about OGT’s function in the blastocyst using this system.

In conclusion, indirect transcriptomic evidence shows that in preimplantation embryos the AID degron induces OGT depletion, but the O-GlcNAc perturbation is suboptimal and necessitates alternative perturbation strategies in this *ex vivo* model. It is worth mentioning that recently optimized versions of the AID system might achieve a better result in preimplantation embryos (Yesbolatova et al, 2020).

### Progressive dampening of cellular activities after acute degradation of endogenous OGT *in vitro*

Auxin-inducible OGT degradation proved to be highly efficient in MEFs, hence we leveraged this cell model to gain insights into the immediate transcriptional response after OGT acute depletion and the evolution of this response over time. We performed mRNA-Seq of three different clones of *OsTIR*,*Ogt^AID^* MEFs treated with auxin for 24, 48 and 96 hours and compared them with the same clones grown in parallel in the absence of auxin (Fig. 4D). Because the addition of the AID tag halves OGT protein amount (Fig. 4C and S4C), we also compared *OsTIR*,*Ogt^AID^* versus *OsTIR*,*Ogt^WT^*in the absence of auxin and found about one hundred DEGs (Fig. S6A, left), mainly involved in actin-based cell motility, a typical fibroblasts’ function (Fig. 4E). Additionally, the side effect of auxin was controlled by treating clones devoid of the AID tag (*OsTIR*,*Ogt^WT^*); we found a small but not negligible gene expression change caused by the drug (Fig. S6A, right) and involving sterol metabolism, which was downregulated (Fig. 4E). All DEGs appearing upon auxin addition in control *OsTIR*,*Ogt^WT^* clones were excluded from the gene ontology (GO) analysis of all the other comparisons.

Acute depletion of endogenous OGT significantly affected the expression of hundreds of genes, with most of the log_2_FC below 1 in both directions (Fig. 4D). Among the most significant changes, we found *Oga* transcripts’ downregulation and the reciprocal *Ogt*’s upregulation (Fig. 4D and S6B), as expected from previous literature showing that O-GlcNAc homeostasis is tightly controlled by rapid adaptation of OGT and OGA protein levels (Qian et al, 2018; Willems et al, 2017; Pravata et al, 2019; Vaidyanathan et al, 2017; Park et al, 2017; Zhang et al, 2014). DEGs in *OsTIR*,*Ogt^AID^*cells after auxin treatment at the three time points overlapped significantly (Fisher exact test’s p-value << 1e^-100^ for all comparisons between two time points; Fig. S6C). A few gene ontology terms were enriched among upregulated DEGs and only at the earliest 24 hour time point, indicating an acute response to OGT or O-GlcNAc depletion leading to an increase of transcripts encoding proteasomal subunits and proteins involved in ribosome biogenesis and mitochondrial translation (Fig. 4E and S6D). Instead, numerous terms were consistently downregulated after OGT depletion, with statistical significance increasing from 24 to 96 hours of acute OGT depletion (Fig. 4E and S6D). The most affected cellular functions at all time points were the mesenchymal transcriptional program, including the actomyosin machinery (e.g. *Cnn1*, *Myl9*) and transforming growth factor beta 3 (*Tgfb3*) (Grafe et al, 2017), and intracellular membrane trafficking. Additionally, after 48 hours of OGT acute depletion, proteasomal subunits and proteins involved in ribosome biogenesis started to be downregulated (Fig. 4E and S6D), although other than the ones upregulated at the 24 hour time point. Finally, at the latest time points, genes involved in translation were more broadly affected (Fig. S6D). Notably, it was reported that the loss of non-catalytic functions versus the loss of the O-GlcNAc transferase activity of OGT affects the abundance of different sets of proteins, including the increase in mitochondrial translation-related proteins in the absence of non-catalytic functions and the downregulation of translation-associated factors in the absence of O-GlcNAcylation (Levine et al, 2021). Hence, it can be hypothesized that the transcriptional response observed at the earliest time point has a higher contribution from the abrupt depletion of OGT molecules (Fig. 4C), while the response accumulating over time is due to the following gradual decrease in O-GlcNAc (Fig. S5E).

The gene network analysis showed that downregulated genes were enriched for targets of the transcription factor c-Myc and of mechanistic target of rapamycin kinase 1 (mTORC1), respectively starting from 48 hours and 96 hours of auxin treatment (Fig. 4F). *c-Myc* is a well-studied oncogene stimulating cell growth (Dang, 2013), while mTORC1 senses cellular nutrients’ and growth factors’ availability to balance between anabolism and macromolecule recycling (Valvezan & Manning, 2019). Downregulated c-Myc targets include proteasomal subunits, ribosomal proteins, translation initiation factors and *Hdac2* (Fig. 4F). The dampening of the two growth pathways was accompanied by signs of activation of a p53 stress response (Fig. 4F), although there was no significant enrichment of gene networks associated with apoptosis or cellular senescence. p53 signaling can repress translation via multiple post-translational and transcriptional mechanisms, including interfering with c-Myc’s activity (Marcel et al, 2015), thus the later-appearing downregulation of protein synthesis-related genes could be a consequence of a primary stress response to the lack of OGT. On the other hand, c-Myc has been reported to be O-GlcNAcylated (Chou et al, 1995) and a recent study found that the O-GlcNAcylation of a regulator of mTORC1, known as Raptor, signals glucose status (Xu et al, 2023), thus O-GlcNAc could also act directly on these pathways. Our experiment does not allow establishing if a relationship of causality exists between the stress response and the dampening of anabolic transcription caused by OGT reduction.

In summary, our transcriptional kinetic study in a cellular model, that is not confounded by evolving cell lineages as in the embryo model, revealed that acute OGT depletion results in the progressive dampening of genes involved in basal cellular functions that include intracellular trafficking, the default MEFs’ cell fate transcriptional program and translation.

## Discussion

Several lines of evidence point towards an evolutionary conserved role for intracellular glycosylation in embryonic growth and development, by modulating the proliferative (Capotosti et al, 2011b) and metabolic capacities of the cell (Yi et al, 2012; Li et al, 2023) and the organism as a whole (Ruan et al, 2014; Kim et al, 2017; Love et al, 2010). However, the early lethality of mouse embryos lacking a maternally inherited functional copy of *Ogt* has hampered molecular investigations of O-GlcNAc function in mammals *in vivo*. In this work, we overcome this obstacle by creating a suite of mouse *Ogt*-hypomorphic mutants and by identifying alleles which perturb O-GlcNAc homeostasis while being compatible with embryonic development (and also not causing signs of cellular lethality in the molecular data). Moreover, the point mutations were introduced to OGT’s glycosyltransferase catalytic core and affect the O-GlcNAc transferase activity specifically. In fact, the four substitutions do not have a predictable effect on OGT’s stability based on *in vitro* characterization (Martinez-Fleites et al, 2008) and *in vivo* our observation that the embryonic survival of the resulting mutants mirrors the *in vitro* decrease in OGT’s glycosyltransferase catalytic rate indicates that the observed phenotypes are caused by impaired O-GlcNAcylation. Of note, OGT’s glycosyltransferase catalytic core is also the site for an additional enzymatic activity, which is the proteolytic cleavage of the cell cycle regulator HCF-1 (Capotosti et al, 2011a; Lazarus et al, 2013a), whose maturation is proposed to ensure a proper progression through S phase and cytokinesis (Julien & Herr, 2003). However, while OGT’s O-GlcNAc transferase activity was found to be necessary for the proliferation of cycling cells, it does not seem to be the case for the cleavage of HCF-1 by OGT (Levine et al, 2021). In addition, the H568A mutation (the most severe hypomorphic allele of this study) does not affect HCF-1 proteolysis (Lazarus et al, 2013b).

Therefore, while it has been known for 20 years that maternal *Ogt* is essential for embryonic survival (O’Donnell et al, 2004; Shafi et al, 2000), our allelic series uncovers the requirement of the O-GlcNAc transferase activity of OGT, thus O-GlcNAc homeostasis itself, to embryonic development, independently of potential non-catalytic roles of OGT. The lethality of heterozygous females with the most disruptive mutations additionally reveals that O-GlcNAc’s essential function starts in the oocyte or in the cleavage stage embryo. The developmental requirement of O-GlcNAcylation was not obvious: OGT’s N-terminus is composed by 13 tetratricopeptide (TPR) repeats mediating interactions with other proteins (Allan & Ratajczak, 2011) and it is reported to be part of many cytosolic and chromatin complexes where it could potentially also play an essential structural role.

Then, it is difficult to predict which OGT’s targets are responsible for the lethality of O-GlcNAc-depleted embryos. Due to the necessity of O-GlcNAc for cell proliferation, cell cycle defects could be involved, although we did not observe the activation of cell cycle checkpoints in our transcriptomics data. Recently, mitochondrial dysfunction triggered by mTOR hyperactivation was proposed to be the cause of cellular lethality following *Ogt*-KO in mESCs (Li et al, 2023). We did not find signs of oxidative stress in the transcriptomes of *Ogt-*hypomorphic embryos, however this remains a possibility that would require to be tested through assays of mitochondrial function.

We analyzed the mouse mutants for which we could obtain alive hemizygous male embryos. The transcriptional effect of the most severe perturbation of O-GlcNAc homeostasis (*Ogt*-T931A/del) *in vivo* suggests a mild developmental delay of the preimplantation mutant embryos. The deregulated transcriptome of E12.5 placentae from less severe mutants (*Ogt*-Y851A) also points in the same direction.

It is worth discussing the delayed placental morphogenesis found in *Ogt*-Y851A mutants. The placenta being a key regulator of embryonic growth, it is conceivable that placental dysfunction has a greater contribution than defects in the embryo proper to embryonic lethality caused by milder reductions of O-GlcNAc. The postimplantation sub-lethality of the *Ogt*-Y851A mutants is consistent with a threshold effect lying in insufficient growth support from the placenta. Future studies of mutants with a placental-specific genetic rescue of *Ogt* function should answer this question.

Because of OGT’s double dose in the female extraembryonic ectoderm (Cheng et al, 2019), another consideration is that the ExE and the deriving placenta are good candidates to be responsible for the observed higher peri-/postimplantation mortality of males upon *Ogt*’s dysfunction. Remarkably, the double dose of *Ogt* in female placentae is conserved in humans (Howerton et al, 2013), making further understanding of the sexually-dimorphic role of *Ogt* in this tissue relevant for human fertility and diseases. For example, it would be interesting to investigate the contribution of placental dysfunction to OGT-XLID neurodevelopmental syndrome. Of note, male-specific higher sensitivity to maternal stress due to lower placental OGT dose has been linked to neuronal effects, namely hypothalamic-pituitary-adrenal stress axis hyperresponsivity as well as changes in hypothalamic gene expression (Howerton et al, 2013; Howerton & Bale, 2014).

The transcriptional analysis of a developing mutant embryo at a specific stage, as performed here, is confounded by the accumulation of defects due to O-GlcNAc perturbation at all previous stages. Hence, the consequences of O-GlcNAc depletion in a static system such as primary MEFs can suggest affected cellular functions lying behind the observed developmental phenotype. By analyzing the transcriptome of MEFs in a time course experiment following endogenous OGT degradation using the auxin system, we found that hypo-O-GlcNAcylation causes the dampening of various basal cellular activities, including translation. Due to the numerous *Ogt* targets, it is likely that O-GlcNAc acts on growth through multiple pathways; thus, it is very difficult to pinpoint the exact ones involved in the observed ultimate transcriptional response. Nonetheless, it is worth noticing that some cellular functions are consistently downregulated in the two models (MEFs, embryo): intracellular membrane trafficking as well as actin-related processes. Moreover, O-GlcNAc-depleted embryos show evidence of a stress response involving translation, precisely the increase in stress granules.

The other molecular effect observed in multiple datasets analyzed in this study is the disturbance of chromatin-based gene silencing upon O-GlcNAc perturbation: in blastocysts, we uncovered a low-magnitude upregulation of retrotransposons which are normally silenced during the course of preimplantation development; in placentae, chromatin gene ontology terms are upregulated and this is associated with a mild upregulation of retrotransposons also in this tissue. In this study, we report for the first time that retrotransposons upregulation occurs *in vivo* when O-GlcNAc homeostasis is perturbed, even moderately. The heterochromatin component or silencing machinery responsible for this phenotype remains to be elucidated. Recent work shows that *Ogt*-KO causes a genome-wide low level of DNA-hypomethylation and that the association between OGT and TET proteins is the responsible player (Sepulveda et al, 2024), but other possibilities are conceivable and need further testing.

Our comparative study of different degrees of impairment of OGT’s activity set the thresholds for maternal inheritance and hemizygous embryonic lethality. We found that the *Ogt^Q449N^* allele, which drastically reduces the enzymatic activity to 2% of the WT, causes lethality to both males and females when it is maternally inherited. In contrast, the milder reduction of OGT’s enzymatic activity to 18% of that of the WT caused by the *Ogt*-T931A substitution resulted in low frequency of maternal transmission of this allele and was embryonic lethal for hemizygous. The mildest reduction of OGT’s activity tested (*Ogt*-Y851A, 24% as compared to WT) is compatible with maternal inheritance and life for both homozygous and hemizygous. To our knowledge, Y851A is the second reported murine *Ogt* mutation that perturbs O-GlcNAc homeostasis while being compatible with adult life and reproduction, besides the very recently described murine *Ogt*-C921Y, which is found in OGT-XLID patients. Based on structural data, the role of C921 in OGT’s catalytic mechanism is more difficult to predict than for the mutations we introduced (Martinez-Fleites et al, 2008). Although the C921Y substitution was shown to reduce OGT V_MAX_ *in vitro* to 25% of the WT (Omelková et al, 2023), these mutant mice do not show embryonic sub-lethality, but present features shared with XLID human patients, making this model very well suited to study XLID’s pathology (Authier et al., 2024). Our *Ogt^Y951A^* mutant should be useful for future investigations of the consequences of prolonged reduction of O-GlcNAcylation in development and adulthood, in physiology and disease mechanisms relevant for human health, such as diabete mellitus (Kim et al, 2017; McClain et al, 2002).

## Materials and Methods

### Animal care and handling

Mice were handled according to the rules and regulations of the EMBL Institutional Animal Care and Use Committee (IACUC) under protocol number 21-012_RM_MB. All procedures involving mice are in compliance with Italian Law (DL 26/2014, EU 63/2010), under protocols number 17/2019-PR to C.L. and 598/2023-PR to M.B. (Ministero della Sanità, Roma, Italy).

Mice were housed in the pathogen-free Animal Care Facility at EMBL Rome on a 12-hours light-dark cycle in temperature and humidity-controlled conditions with *ad libitum* access to food and water.

Generation of murine *Ogt* allelic series: *Ogt^Y851A^, Ogt^T931A^, Ogt^Q949N^, Ogt^H568A^, Ogt^NLS-^, Ogt^NterAID-MYC-FLAG^ Ogt* mutant alleles were created by CRISPR/Cas9-editing in the zygote (FVB/NCrl genetic background, Charles River) as previously described (Quadros et al, 2017). Transgenic mouse production was performed by the Gene Editing and Virus Facility at EMBL Rome. Sequences of the single-stranded donor templates are provided in Table S1.

Briefly, for *Ogt^Y851A^* (FVB/NCrl-*Ogt*^em1*(Y851A)*^Emr) CRISPR crRNA oligo (TAGATGGGACGTCTACACCC) was annealed with tracrRNA and combined with a homology flanked ssODN donor coding for Tyrosine at amino acid position 851, substituting the wild type Alanine. The target location was exon 19 of transcript *Ogt-201*, (ENSEMBL v109); genomic coordinate: ChrX. 100719847-100719886 (GRCm39/mm39).

Similarly, for *Ogt^T931A^* (FVB/NCrl-*Ogt*^em2(*T931A*)^Emr) crRNA oligo (TTGTGTAATGGACACACCAC) was combined with a ssODN donor coding for Threonine at amino acid position 931 substituting Alanine, exon 20, transcript *Ogt-201*, ChrX. 100722515-100722518.

For *Ogt^Q949N^* (FVB/NCrl-*Ogt*^em3(*Q949N*)^Emr) crRNA oligo (TAGATGGGACGTCTACACCC) was combined with a ssODN donor coding for Glutamine at amino acid position 949 substituting Asparagine, exon 19, transcript *Ogt-201*, ChrX. 100719847-100719886.

For *Ogt^H568A^* (FVB/NCrl-*Ogt*^em4(*H568A*)^Emr) crRNA oligos (GCTATGTGAGTTCTGACTTC and ATGAAGTGTGGAATACGTCA) were combined with a ssODN donor coding for Histidine at amino acid position 568 substituting Alanine, exon 13, transcript *Ogt-201*, ChrX. 100713458->100713464.

For *Ogt^NLS-^* (FVB/NCrl-*Ogt*^em5(*DFPmut*)^Emr) crRNA oligos (CAATAAGCATCAGGAAAGTC and GCCAAGTTACAATAAGCATC) were combined with a ssODN donor coding for three Alanines at amino acid positions 461-463 substituting three amino acids associated with *Ogt* NLS, Aspartic acid, Phenylalanine and Proline, exon 11, transcript *Ogt-201*, ChrX. 100713458->100713464.

For *Ogt^NterAID-MYC-FLAG^* (FVB/NCrl-*Ogt*^em6(*AID*)^Emr) crRNA oligos (CTCCAGATGGCGTCTTCCGT and ACTGTCGGCCACGTTGCCCA) were combined with a ssDNA donor coding for AID, 2xMyc and 1xFLAG tags inserted at *Ogt* N-terminus, transcript *Ogt-201*, ChrX. 100683892.

In all cases, annealed sgRNAs were complexed with Cas9 protein and combined with their respective ssDNA donors (Cas9 protein 50 ng/µL, sgRNA 20 ng/µL, ssDNA 20 ng/µL. All single stranded DNA and RNA oligos were synthesized by IDT. Cas9 protein (IDT), sgRNA and ssDNA donor template were co-microinjected into zygote pronuclei using standard protocols (Liu & Du, 2019) and after overnight culture 2-cell embryos were surgically implanted into the oviduct of day 0.5 post-coitum pseudopregnant CD1 mice.

Founder mice were screened for the presence of the mutation by PCR, first using primers flanking the sgRNA cut sites which identify InDels generated by NHEJ repair and can also detect larger products implying HDR. Secondly, 5’ and 3’ PCRs using the same primers in combination with template-specific primers allowed for the identification of potential founders. These PCR products were then Sanger sequenced and aligned with the *in silico* design.

The *Ogt* mutant strains were maintained on an FVB/NCrl genetic background. PCR genotyping using gDNA isolated from tail biopsies was performed at weaning (postnatal day 21), using primers listed in Table S2.

### Generation of the *ROSA26^OsTIR^* allele

The *OsTIR1* cDNA was modified by the addition of Myc and HA tags for protein detection (Nishimura et al, 2009). The *OsTIR1-Myc-HA* transgene was next inserted in a targeting vector specific to the ROSA26 locus. A *lox-stop-lox* (*LSL*) cassette was added before the transgene to prevent protein expression. The *LSL-OsTIR1* targeting construct was electroporated into A9 ESCs (129xC57BL/6 genetic background) (Fazio et al, 2011). Southern blotting of the individual ESC-clone-derived DNA was used to identify homologous recombinants. A9 ESCs with successful insertion were injected into C57BL/6 8-cell stage embryos that were then transferred into foster mothers as previously described (Fazio et al, 2011). The resulting chimeras were tested for germline transmission. F_1_ mice bearing the *OsTIR1-Myc-HA* transgene were crossed to the FLP-expressing transgenic mice (FLPeR) (Farley et al, 2000) to remove the frt-flanked neo cassette. The resulting progeny was crossed for 12 generations with C57BL/6N animals to obtain the C57BL/6N-ROSA26^tm1.1*(osTIR1)*^ChLan mouse line (ROSA26-*LSL-OsTIR1*). To remove the *LSL* cassette, the ROSA26-*LSL-OsTIR1* mice were crossed to the Cre-Deleter strain (Schwenk et al, 1995), generating the FVB;B6J;129-Gt(ROSA)26Sor^tm1*(OsTIR)*^Emr allele (hereafter ROSA26*^OsTIR^*), constitutively expressing *OsTIR1*. The ROSA26*^OsTIR^* allele was then backcrossed on FVB/NCrl genetic background for 2 generations.

### Dissection of mouse placentae

Embryos were dissected at E12.5 post-coitum from the uteri of naturally mated mice. Placentae were dissected in phosphate-buffered saline solution (PBS) + 0.1% Bovine Serum Albumin (BSA) and the exterior decidua was peeled out in order to minimize the presence of maternal tissue. For RNA-Seq, clean placentae were bisected via transverse sectioning; one half was collected in a 1.5 mL tube containing 300 µL of 1x RNA Protection Reagent (NEB #T2011-1) and snap-frozen. The E12.5 whole embryos were cleaned from the yolk sac and imaged with trans-illumination on Leica M205C dissection microscope. For limb staging, the images were analyzed using the eMOSS software (https://limbstaging.embl.es/) (Musy et al, 2018).

### Derivation and culture of mouse embryonic fibroblasts (MEFs)

Primary MEFs were derived from E12.5 post-coitum embryos as previously described (Jain et al, 2014). Briefly, the head and the organs were removed, and gDNA was extracted from the head using the PCRBIO Rapid Extract Lysis Kit (PCR Biosystems #PB15.11-24) for PCR genotyping with primers described in Table S3. The embryonic body was minced using sterile forceps and small pieces of tissue were transferred to 5 mL Trypsin-EDTA (0.05%; Gibco #25300054) and homogenized through repeated pipetting. The homogenized tissue was spun down, the supernatant was removed and the dissociated cells resuspended in MEF medium (Dulbecco’s Modified Eagle Medium (DMEM; Gibco #41965-039), 10% fetal bovine serum (PANBiotech #3306-P131004), 100 U/mL Penicillin/Streptomycin (Gibco #15140122), 1x GlutaMAX (Gibco #35050061)). Cells were cultured on 0.1% gelatin in an incubator at 5% CO_2_ and 37°C. MEFs were passaged at ∼80% confluency using Trypsin-EDTA (0.05%) for cell detachment and dissociation. All experiments were performed between passage 1 and 4. The three pairs of clones studied were derived from three different litters, each pair (*OsTIR*,*Ogt^AID^*and control *OsTIR*,*Ogt^WT^*) coming from the same litter.

### Auxin treatment of MEFs

Indole-3-acetic acid sodium salt, or IAA or auxin (Sigma-Aldrich #I5148), was resuspended in ddH_2_O under sterile conditions to a stock concentration of 10 mM, aliquoted and stored at −20 °C. For auxin treatment of MEFs, 500 µM of auxin was added to the culture medium at day 0 and cells were collected 24, 48 and 96 hours later. For the 96-hour time point, the medium was changed after 48 hours with fresh MEF medium supplemented with auxin 500 µM, in order to keep auxin concentration constant throughout the experimental time. At the time of collection, 500 µM auxin was added to all reagents used for cell dissociation (Trypsin-EDTA and DPBS). Cells were collected in 1.5 mL tubes, resuspended in 500 µL of DPBS (Gibco #14190-094) with or w/o 500 µM auxin and the same volume of 2x Monarch DNA/RNA Protection Reagent (NEB #T2011-1) was added before snap-freezing and storage at −80°C until RNA extraction.

### Collection of embryos from natural mating

Superovulation of 6-12 weeks-old FVB females was induced by hormonal stimulation (5 IU of PMSG and 5 IU of hCG 64 hr and 16 hr before collection, respectively) and cumulus-oocyte complexes were collected in warm M2 medium (Sigma-Aldrich #M7167), then moved to a drop of warm hyaluronidase (Sigma-Aldrich #H4272-30 mg). Individual oocytes were collected, washed 4-5 times in M2 and then moved to culture.

### In vitro fertilization

In vitro fertilization (IVF) was performed based on the published protocol, with minor modifications (Guan et al, 2014). When WT sperm was used for IVF experiments, the strain was always FVB/NCrl except for the *Ogt^T931A^* blastocysts Smart-Seq experiment where it was PWD/PhjEmr. Superovulation of 6-12 weeks-old FVB females was induced by hormonal stimulation (5 IU of PMSG (Prospec Bio #HOR-272) and 5 IU of hCG (Sigma-Aldrich #CG5-1VL) 64 hours and 16 hours before collection, respectively), and cumulus-oocyte complexes were collected in KSOM medium containing GSH (final concentration 10 mM; Sigma-Aldrich #G6013). Concomitantly, cauda epididymis and vasa deferentia from FVB or PWD males were dissected and sperm was gently squeezed out into capacitation medium (HTF supplemented with MBCS at final concentration of 0.75 mM; HTF: Sigma-Aldrich #MR-070-D, MBCD: Sigma-Aldrich #C4555) and allowed to swim up for 1 hour. Sperm were subsequently counted and cumulus-oocyte complexes were inseminated with 0.2 million sperm in a 200 µL fertilization drop. Four hours after sperm addition, zygotes were cleaned from the surrounding cumulus cells and sperm by 5-6 washes in KSOM, then transferred to culture media. Embryos were cultured in a standard mammalian cell incubator (37 °C, 5% CO_2_) in EmbryoMax KSOM Mouse Embryo Medium (Sigma-Aldrich #MR-106-D).

### Auxin treatment of embryos grown *ex vivo*

For auxin treatment of zygotes derived from IVF, 250 µM of auxin (and 500 µM for OGT staining 3.5 hours post-IVF) was added to the fertilization drop (KSOM medium containing GSH), to all KSOM washing drops and to the KSOM medium used for culture for the next following days. In order to maintain a constant auxin concentration, embryos were moved to fresh KSOM supplemented with 250 µM auxin every second day. For auxin treatment of the morulae, 72 hours after collection from natural mating, embryos were moved to fresh media either not containing or containing 250 µM of auxin for culture for the next 24 hours.

On the day of fixation for immunofluorescence staining or collection for Smart-Seq, M2 medium was supplemented with 250 µM (or 500 µM) auxin. For Smart-Seq, groups of 5-10 blastocysts were washed twice in warm M2 drops and collected in 4 uL of a lysis buffer containing 0.5% Triton X-100 in H_2_O, 1 U/μL SUPERase•In RNase Inhibitor (ThermoFisher #AM2696), 2.5 mM dNTPs (BiotechRabbit #BR0600204, 10 mM each) and 2.5 μM oligo-dT primer (5′–AAGCAGTGGTATCAACGCAGAGTACT30VN-3′). Blastocysts were collected in five different replicates of breeding followed by auxin treatment and stored at −80 °C prior to Smart-Seq library preparation and next generation sequencing.

### Immunofluorescence staining of preimplantation embryos

Zygotes (3.5 hours post-IVF) and morulae (70 hours post-IVF) were treated the same way, as previously described in Bošković *et al*. (Bošković et al, 2012) with some minor modifications as follows. After two washes in M2 medium (Sigma-Aldrich #M7167), the zona pellucida was removed by a brief incubation in drops of warm Acidic Tyrode’s solution (Sigma-Aldrich #MR-004-D), followed by other two M2 washes to neutralize the acid. The embryos were then washed once in PBS + 0.1% BSA (Sigma-Aldrich #A2153), fixed in 4% paraformaldehyde (PFA) in PBS for 20’ at 37 °C, permeabilized in 0.5% Triton X-100 for 20’ at 37 °C and washed three times in PBS-T (0.15% Tween-20 in PBS). The epitope was then unmasked in 50 mM NH_4_Cl solution in H_2_O for 10’ at room temperature, followed by two additional PBS-T washes, and then blocked for 3 hr at room temperature or overnight at 4 °C in BSA (Sigma-Aldrich A2153) 3% in PBS-T. Primary antibody incubation was performed overnight at 4 °C in the blocking solution, followed by three washes in PBS-T, re-blocking for 30’ at room temperature, three additional PBS-T washes, secondary antibody incubation for 1 to 2 hr at room temperature in the blocking solution and three final PBS-T washes. The embryos were immediately mounted on coverslips in Vectashield (Vector Laboratories #H-1200) containing 4’-6-Diamidino-2-phenylindole (DAPI) or, in the case of the morulae, in drops of 75% Vectashield in PBS in order to preserve the 3D structure. Fixed immunostained samples were imaged using a Nikon AX scanning confocal (using galvanometric mirrors).

The primary antibodies used were: anti-OGT (Abcam #ab177941), anti-O-GlcNAc clone RL2 (Abcam #ab2739 and Merck Millipore #MABS157). Dilution of all primary antibodies was 1:200. Secondary antibodies used were: A647-conjugated goat anti-rabbit IgG (ThermoFisher #A21244) and A647-conjugated goat anti-mouse IgG (ThermoFisher #A21236). Dilution of all secondary antibodies was 1:500.

### Collection of *Ogt^T931A^* blastocysts and RNA extraction

92 hours post-IVF, groups of 5-10 embryos from *Ogt^T931A/+^* or *Ogt^+/+^*/*Ogt^T931del/+^* mothers were washed twice in warm M2 drops. Single blastocysts were collected from the M2 drop in 5 µL of 1x TCL lysis buffer (Qiagen #1031586) containing 1% (v/v) 2-mercaptoethanol (Gibco #31350010) and 0.5 U/µL of SUPERase•In RNase Inhibitor (ThermoFisher #AM2694).

Total RNA was purified using RNAClean XP beads (Beckman Coulter #A63987) according to the manufacturer’s protocol for a 96-well plate and Small Volume Reactions. After the last ethanol wash, the RNA was eluted from RNA beads in 8 µL of H_2_O, 3 µL of which were transferred to a new 96-well plate containing 1 µL of dNTPs (BiotechRabbit #BR0600204, 10 mM each) and 1 µL of 10 µM oligo-dT primer (5′–AAGCAGTGGTATCAACGCAGAGTACT30VN-3′). The plate was stored at −20 °C prior to Smart-Seq library preparation and next generation sequencing.

Single blastocysts from *Ogt^T931A/+^* mothers were genotyped using cDNA. The cDNA from single blastocysts was prepared from 3 μL of RNA (extracted as described above) using SuperScrip IV RT (ThermoFisher #18090200) according to the manufacturer’s instructions and using random hexamers. PCR genotyping of the resulting cDNA was performed with primers in Table S4 and the result was confirmed with Sanger sequencing of the PCR product.

### Smart-Seq library preparation and sequencing from single embryo

Single-embryo full-length cDNA libraries for mRNA sequencing were prepared by EMBL Genomics Core Facility using a modified Smart-Seq2 protocol (Picelli et al, 2014) utilizing SuperScript IV Reverse Transcriptase (ThermoFisher #18090200) and the tagmentation procedure previously described (Hennig et al, 2018). The retrotranscription reaction mix was as follows: 2 μL SSRT IV 5x buffer, 0.5 μL 100 mM DTT, 2 μL 5 M betaine, 0.1 μL 1 M MgCl_2_, 0.25 μL 40 U/μL RNAse inhibitor (Takara #2313A), 0.25 μL SSRT IV, 0.1 μL 100 µM TSO, 1.15 μL RNase-free H_2_O; with thermal conditions: 52 °C 15’, 80 °C 10’. cDNA was amplified using 18 PCR cycles. The cDNA cleanup (0.6x SPRI beads ratio; Beckman Coulter #B23319) was carried out omitting the ethanol wash steps and the elution volume was 13 μL of H_2_O. For tagmentation, the sample input was normalized to 0.2 ng/μL. Before the final clean-up after tagmentation and PCR, 2 μL of each sample were pooled in a single tube. The pool was cleaned-up using 0.7x SPRI ratio and sequenced in one run (40 bp paired-end mode for *Ogt^T931A^* and 75 bp single-end for *Ogt^AID^*) on the Illumina NextSeq500 sequencer. Table S5 contains the number of pooled embryos and average number of reads obtained per embryo for each library.

### MEFs mRNA-Seq library preparation and sequencing

Total RNA was extracted with Monarch Total RNA Miniprep Kit (NEB #T2010S). The integrity of the RNA was determined using the TapeStation 4150 (High Sensitivity RNA kit, Agilent Technologies #5067-5579). Library preparation was performed by EMBL Genomics Core Facility using the TruSeq Stranded mRNA Library Preparation kit according to the manufacturer’s instructions (Illumina #20020594). All libraries were pooled and sequenced in one run (75 bp single-end mode) on the Illumina NextSeq500 sequencer at the Genomics Core Facility of EMBL Heidelberg.

### mRNA-Seq library preparation and sequencing from single placentae

Samples were thawed and homogenized using BioSpec 11079110Z Zirconia/Silica Bead 1.0 mm Diameter (3 cycles, 90 s each, 2500 rpm), supernatant was collected in a new tube and treated 10’ at 55 °C with Proteinase K (15 μL of enzyme every 300 μL of sample, provided with Monarch Total RNA Miniprep Kit). Debris were removed by centrifugation at 16000 x g for 2’ and collection of the supernatant. RNA was then extracted by adding an equal volume of RNA lysis buffer and proceeding with Monarch Total RNA Miniprep Kit (NEB #T2010S) extraction, according to the manufacturer’s instructions. After RNA’s elution with water, an additional treatment with 5U/50μL Turbo DNAse (Thermo Scientific #AM2238) was performed for 15’ at 37 °C in order to reduce DNA contamination for RNA repeats analysis. The RNA samples were cleaned with Monarch RNA Cleanup Kit (NEB #T2030) according to the manufacturer’s instructions. The integrity of the RNA was determined using the TapeStation 4150 (High Sensitivity RNA kit, Agilent Technologies #5067-5579). Library preparation was performed by EMBL Genomics Core Facility using the NEBNext Ultra II Directional RNA Library Prep Kit for Illumina (NEB #E7760L), in combination with the NEBNext Poly(A) mRNA Magnetic Isolation Module for mRNA enrichment (NEB #E7490L), according to the manufacturer’s instructions. All libraries were pooled and sequenced in one run (40 bp paired-end mode) on the Illumina NextSeq500 sequencer at the Genomics Core Facility of EMBL Heidelberg.

### Cellular fractionation

About 10 million cells were collected, washed with PBS and resuspended in 10 volumes of hypotonic buffer (10 mM HEPES pH 7.65; 1.5 mM MgCl_2_, 10 mM KCl, 0.5 mM DTT (Invitrogen #15508-013)). Cells were then incubated for 15’ at 4 °C under gentle agitation and Dounce homogenized 20 times using a tight pestle. The homogenized nuclei were centrifuged at 228 x g for 5 min at 4 °C. The supernatant (cytosol) was clarified by high-speed centrifugation (5 min, 20000 x g, 4 °C). The nuclei were washed twice in 2 ml of washing buffer (15 mM HEPES pH 7.65; 10 mM MgCl_2_, 250 mM sucrose, 0.5 mM DTT). Nuclei were lysed by resuspension in RIPA buffer (150 mM NaCl, 1% NP-40, 0.5% deoxycholic acid, 0.1% SDS, 50 mM Tris pH 8.0 in ddH_2_O).

### Western blotting

Total protein lysates were prepared by lysing cells with RIPA buffer and subsequent high-speed centrifugation (5 min, 20000 x g, 4 °C) to eliminate insoluble fractions. Protein concentration was quantified using the Pierce BCA Protein Quantification kit (ThermoFisher #23225). 20 μg of protein extract and 5 μL of WesternSure Pre-stained Chemiluminescent Protein Ladder (Li-cor #926-98000) were loaded in a NuPAGE 4-12% Bis-Tris or Novex 4-20% Tris-Glycine Protein Gel, 1.0 mm (ThermoFisher #NP0335BOX and #XP04200BOX) and run at 150V in an XCell Sure Lock apparatus (ThermoFisher). Proteins were transferred on a PVDF membrane in a Mini-Protean Tetra System apparatus or a Trans-Blot Turbo Transfer System (Bio-Rad). Abundant epitopes were blocked with a buffer containing 5% milk or, for O-GlcNac detection, 5% BSA (Sigma-Aldrich #A2153), and 0.1% Tween-20 in ddH_2_O. Membranes were incubated overnight at 4 °C (or for 1 hour at room temperature for loading controls) with primary antibody dilutions in a buffer containing 5% BSA and 0.1% Tween-20 in ddH_2_O. Membranes were washed twice with 0.5% Triton X-100 and 0.5 M NaCl in ddH_2_O for 5’ and once with 0.5 M NaCl in ddH_2_O for 10’, then rinsed with PBS and incubated for 1 hour at room temperature with 1:200000 secondary antibody dilutions of HRP-coupled goat anti-rabbit IgG (ThermoFisher #G-21234) or 1:100000 dilutions of HRP-coupled goat anti-mouse IgG (ThermoFisher #G-21040). Membranes were washed as after primary antibody and incubated with GE Healthcare ECL Prime (Sigma #RPN2232) for imaging using a ChemiDoc imaging system (BioRad) or ImageQuant 800 (Amersham).

The primary antibodies used were: anti-OGT (Abcam #ab177941) 1:2000, anti-O-GlcNAc clone RL2 1:2000 (Abcam #ab2739 and Merck Millipore #MABS157), anti-lamin A/C (Santa Cruz Biotechnology #sc-376248) 1:200, anti-histone H3 1:100000 (Abcam #ab1791), anti-vinculin 1:5000 (Sigma-Aldrich #V9264).

### Cell immunofluorescence

MEFs were seeded a day before fixation in 12-well plates on coverslips covered with 0.1% gelatin. Cells were fixed for 10’ at room temperature using PFA/SEM buffer (4% PFA, 0.12 M sucrose, 3 mM EGTA, 2 mM MgCl_2_ in PBS), then washed once with PBS. Aldehyde groups were quenched with 50 mM ammonium for 10’, then cells were washed once with PBS and permeabilized with 0.1% Triton X-100 for 10’. Cells were washed three times with PBS, then unspecific epitopes were saturated using 5% goat serum for 30’ at room temperature. Coverslips were incubated overnight at 4 °C with primary antibody anti-OGT (Abcam #ab177941) 1:200 in 1% BSA (or with 1% BSA w/o primary antibody, as a control), the day after washed three times with PBS and incubated with the secondary antibody (A488-conjugated goat anti-rabbit IgG (ThermoFisher #A11008)) diluted 1:1000 in 1% BSA for 30’ at room temperature. Coverslips were washed once with PBS, incubated with DAPI 1:1000 in PBS for 5’ at room temperature and washed again with PBS. Coverslips were mounted on glass slides in ProLong Diamond Antifade Mountant (ThermoFisher #P36961). Fixed immunostained samples were imaged on a Nikon A1 confocal microscope.

### Quantification of microscopy images

All steps were performed using ImageJ (Schneider et al, 2012) when not differently specified. For the quantification of O-GlcNAc signal in single morulae (Figure S5F): i. brightness and contrast of the raw images were adjusted to the same scale, ii. projection of the z stacks was performed for each image using the maximum signal intensity, iii. images were denoised using Aydin (https://doi.org/10.5281/zenodo.5654826), with Classic Image Denoiser butterworth and default parameters, iv. ellipses were drawn inside each embryo and mean intensity in the area was measured for single embryos.

### Analysis of mRNA-sequencing in MEFs and placentae for single copy genes

The analysis pipeline was performed using Galaxy (Afgan et al, 2018) to obtain the table of gene counts. The quality of the reads was analyzed using FastQC v0.11.8 (https://www.bioinformatics.babraham.ac.uk/projects/fastqc/), then reads trimmed from adapters and low-quality 3’-end nucleotides using Trim Galore v0.6.3 (https://www.bioinformatics.babraham.ac.uk/projects/trim_galore/) with default parameters for single-end or paired-end libraries (-q 20 --stringency 1 -e 0.1 --length 20 –paired). Reads were mapped to Gencode vM25 (GRCm38.p6) transcript sequences using Salmon v0.8.2 (Patro et al, 2017) with default parameters for stranded single-end or for paired-end libraries. For MEFs, the *AID-2xMyc-FLAG* tag and the *OsTIR-Myc-HA* transgene had been added to the transcriptome. The gene counts were used in a custom Rmd script as input for DESeq2 v1.34.0 (Love et al, 2014) after gene-level summarization using tximport (Soneson et al, 2016). The test used for statistical significance was the Wald test, the obtained p-values were corrected for multiple testing using the Benjamini and Hochberg method (default in DESeq2) and the significance cutoff for optimizing the independent filtering was 0.05. Whenever MA-plots are shown, the log_2_ fold changes are shrunken using the ‘ashr’ method (Stephens, 2017) and the x-axis shows mean DESeq2-normalized gene counts across samples.

### Analysis of single embryo mRNA-sequencing for single copy genes

The analysis pipeline was performed using Galaxy (Afgan et al, 2018) to obtain the table of gene counts. Briefly, the quality of reads was analyzed using FastQC v0.11.8 (https://www.bioinformatics.babraham.ac.uk/projects/fastqc/). Reads were trimmed from adapters and low-quality 3’-end nucleotides using Trim Galore v0.6.3 (https://www.bioinformatics.babraham.ac.uk/projects/trim_galore/) with default parameters for single-end or paired-end libraries (-q 20 --stringency 1 -e 0.1 --length 20 –paired), before mapping them to the GRCm38 mouse genome using STAR v2.7.8a (Dobin et al, 2013) and default parameters for single-end or paired-end reads. For the *Ogt^AID^* blastocysts, the *AID-2xMyc-FLAG* tag and the *OsTIR-Myc-HA* transgene had been concatenated to the genome fasta and GTF annotation. Gene counts were obtained from the bam files using featureCounts (subreads v2.0.1) (Liao et al, 2014) with default parameters for single-end or paired-end reads and counting fragments instead of reads in the latter case. The gene counts were used in a custom Rmd script as input for DESeq2 v1.34.0 (Love et al, 2014). The test used for statistical significance was the Wald test, the obtained p-values were corrected for multiple testing using the Benjamini and Hochberg method (default in DESeq2) and the significance cutoff for optimizing the independent filtering was 0.05.

For the *Ogt^AID^* blastocysts, pooled from more than one embryo generation and collection experiment, batch effect had to be considered when testing for differential expression using ‘DESeq’ function. To this aim, the package RUVSeq (Risso et al, 2014) was used to compute the factors of unwanted variation, with function ‘RUVg’ and k=6. The subset of control genes to use in RUVg was found with a first run of DESeq2 differential expression analysis, separately for all untreated and all auxin-treated blastocysts: the genes with i. mean DESeq2-normalized counts across samples > 100, ii. adjusted p-value > 0.8 and iii. log_2_FC < 0.05 for each condition were selected. The 6 factors of unwanted variation computed with RUVSeq were used in ‘DESeq’ formula *∼ W1 + … + W6 + genotype*.

For MA-plots, the log_2_ fold changes are shrunken using the ‘ashr’ method (Stephens, 2017) and the x-axis shows mean DESeq2-normalized gene counts across samples. Before principal component analysis (PCA) and differential expression analysis, low-quality samples were removed by discarding embryos with < 10^6^ reads and outlier embryos in a scatter plot of mitochondrial DNA (mtDNA) gene expression versus percentage of reads mapping to ribosomal DNA (rDNA). The exact number of embryos filtered at each step and the final number used for DE analysis is in Table S4.

### *In silico* genotyping of single embryos from mRNA-sequencing data

Sex was assigned to the single embryo transcriptomes based on the DESeq2-normalized counts of chrY-mapping genes *Ddx3y* and *Eif2s3y* (Groff et al, 2019). The assignment of the *Ogt^AID^* genotype was based on the raw sum of reads mapping to the *AID-2xMyc-FLAG* exogenous sequence, whose distribution divided the samples in two distinct populations. A sample was defined as wild type if the sum of *AID-2xMyc-FLAG*-mapping reads was < 72.

### Analyses of gene set enrichment from mRNA-sequencing data

All performed in a custom Rmd script. Gene ontology (GO) over-representation test in MEFs was performed on differentially expressed genes (DEGs) (adj. p-value < 0.05, any log_2_FC) with mean DESeq2-normalized counts across samples > 10 using function ‘enrichGO’ of R package clusterProfiler v4.2.2 (Wu et al, 2021) with adjusted p-value and q-value cutoffs of 0.05 and 0.1, respectively, and default parameters. GO level 5 was then selected using function ‘gofilter’ and results were simplified by q-value using function ‘simplify’ and a similarity cutoff of 0.6. for MEFs) or 0.7 (for blastocysts from the *Ogt^T931A^* allele’s IVF experiment).

Analysis of over-representation of Molecular Signature Database (MsigDb) hallmark gene sets was performed on DEGs with mean DESeq2-normalized counts across samples > 10 using function ‘GSEA’ of R package clusterProfiler v4.2.2 with adjusted p-value cutoff of 0.05 and default parameters.

Gene set enrichment analysis (GSEA) was performed on all genes of the dataset with mean DESeq2-normalized counts across samples > 10, ranked by −log10(adj. p-value)*sign(log_2_FC), using function ‘gseGO’ (for GO terms) or GSEA (for placental markers) of R package ‘clusterProfiler’ v4.2.2, with adjusted p-value cutoff of 0.05 and default parameters. For GO terms, GSEA results were simplified based on adj. p-value after computing semantic similarity using ‘mgoSim’ function of R package ‘GoSemSim’ (Yu et al, 2010). Similarity cutoff was 0.6 for all figures showing GSEA results with GO terms.

### Principal component analysis of single embryo mRNA-sequencing data

The PCAs shown in Figures 2C, S2B,C and S5H were produced with function ‘prcomp’ on log_2_-transformed, DESeq2-normalized counts, using the first 1000 genes with the highest variance, after removing genes with mean DESeq2-normalized counts across samples ≤ 10.

### Analysis of retrotransposons’ expression

The analysis was performed using a custom Snakemake v5.9.1 pipeline (Mölder et al, 2021), available at https://github.com/boulardlab/Ogt_mouse_models_Formichetti2024/blob/master/OgtY851A_placenta_mRNASeq/snake-make/.

In summary, the quality of the fastq files was checked with FastQC v0.11.8 and reads were trimmed using Trim Galore v0.6.4 with default parameters for paired-end libraries (-q 20 --stringency 1 -e 0.1 --length 20 –paired). Trimmed reads were aligned to GRCm38 mouse genome with STAR v2.7.5c (Dobin et al, 2013), with parameters recommended by Teissandier et al. (Teissandier et al, 2019) for the analysis of transcripts derived RNA TEs in the mouse genome: --outFilterMultimapNmax 5000 --outSAMmultNmax 1 --outFilterMismatchNmax 3 --outMultimapperOrder Random --winAnchorMultimapNmax 5000 --alignEndsType EndToEnd --alignIntronMax 1 --alignMatesGapMax 350 –seedSearchStartLmax 30 –alignTranscriptsPerReadNmax 30000 –alignWindowsPerReadNmax 30000 –alignTranscriptsPerWindowNmax 300 -seedPerReadNmax 3000 --seedPerWindowNmax 300 --seedNoneLociPerWindow 1000. After mapping, using a custom script, reads were kept only from pairs where both mates were: - completely included into a repetitive element; - not overlapping with gene bodies of Gencode vM25. Repeat Library 20140131 (mm10, Dec 2011) was used as repetitive elements annotation, after excluding “Simple repeats” and “Low complexity repeats”.

With the STAR parameters above, only one random alignment (that with the highest alignment score) is reported for multimappers, preventing the precise quantification of single repetitive elements. Therefore, for each repName of the repetitive element annotation - often present in multiple copies in the genome - read counts were summarized using featureCounts (subreads v2.0.1) (Liao et al, 2014), with parameters -p -B -s 0 --fracOverlap 1 -M.

The downstream analysis of TEs expression was performed in a custom Rmd script. First, for Fig. 2G and S2F, an analysis at the family level (repFamily field in the Repeat Library annotation) was performed. For this analysis, Fragments per Kilobase per Million (FPKM) were computed for each repFamily by: i. summing counts for all elements belonging to that repFamily; ii. dividing this sum by the number of STAR input reads for that sample (/2 because of featureCounts quantification of fragments instead of single reads) and multiplying by 10^6^; iii. dividing what obtained by the total sum of featureCounts-summarized lengths (in Kb) of all elements belonging to that repFamily. FPKM values of DNA transposon families were manually checked to remove samples with high DNA contamination. Differential Expression analysis of RNA TEs was performed at the repName level, using DESeq2 v1.34.0 (Love et al, 2014) and adding the value of the total sum of DNA FPKM as a confounding factor to DESeq formula (*∼ DNA_FPKM + condition*). The test used for statistical significance was the Wald test, the obtained p-values were corrected for multiple testing using the Benjamini and Hochberg method (default in DESeq2) and the significance cutoff for optimizing the independent filtering was 0.05. For MA-plots, the log_2_ fold changes are shrunken using the ‘ashr’ method (Stephens, 2017) and the x-axis shows mean DESeq2-normalized counts. For Figures S2G and S3D, FPKM values for each repName were computed as follows: i. repName counts (from featureCounts summarization) were divided by the total number of STAR input reads for that sample (/2 because of featureCounts quantification of fragments instead of single reads) and multiplied by 10⁶; ii. what obtained was divided by the featureCounts-summarized length (in Kb) of that repName.

### Analysis of publicly available mRNA-Seq data

The analysis of datasets GSE66582 (Wu et al., 2016) and GSE76505 (Zhang et al., 2017;) for Figures 2F, S2E, S2H was performed as described in (Formichetti et al, 2024).

### Statistical analyses

All statistical tests used are specified in the figure legends. The exact p-values are indicated in the figures, while adjusted p-values are indicated in figures only when <0.05 for single copy genes or placental retrotransposons or <0.1 for preimplantation embryos’s retrotransposons and higher values are considered not significant. For boxplots, hinges correspond to first and third quartiles; median is shown inside; whiskers extend to the largest and smallest values no further than 1.5 * IQR from the hinge (IQR = inter-quartile range, or distance between the first and third quartiles); data beyond the end of the whiskers are plotted individually.

## Data availability

All raw RNA-Seq data generated in this study have been deposited in Biostudies and are available under the following accession codes:

- E-MTAB-13298: "mRNA-Seq of primary MEFs untreated or treated with auxin for OGT degradation" Link for reviewers: https://www.ebi.ac.uk/biostudies/arrayexpress/studies/E-MTAB-13298?key=ceafd6c8-154a-4645-bb8a-907a68126419
- E-MTAB-13297: "SMART-Seq of untreated and auxin-treated blastocysts for OGT degradation" Link for reviewers: https://www.ebi.ac.uk/biostudies/arrayexpress/studies/E-MTAB-13297?key=3a236f18-ae3f-4d75-8969-c602f88e68c2
- E-MTAB-13299: "mRNA-Seq of placentas bearing the OgtY851A mutation" Link for reviewers: https://www.ebi.ac.uk/biostudies/arrayexpress/studies/E-MTAB-13299?key=e2d8ae00-4772-440d-a33b-b1caa54d12e2
- E-MTAB-13499: “Smart-Seq of female and male blastocysts from mothers bearing the OgtT931A and OgtT931del mutations” Link for reviewers: https://www.ebi.ac.uk/biostudies/arrayexpress/studies/E-MTAB-13499?key=4772105a-02c5-4875-9491-ede05d2130f8

## Code availability

The source code of all bioinformatics analyses is available at https://github.com/boulardlab/Ogt_mouse_models_Formichetti2024.

## Acknowledgments

This research was supported by the European Molecular Biology Laboratory (EMBL). We thank members of the Boulard laboratory for helpful discussion and Agnese Loda for critical reading of the manuscript. We thank the EMBL Rome Laboratory Resources Animal Facility and in particular Giuseppe Chiapparelli and Valerio Rossi for their essential support with animal husbandry and genotyping. We acknowledge all members of the Gene Editing and Virus Facility at EMBL Rome for assistance with mouse genome engineering, the EMBL Rome microscopy facility for assistance with microscopy imaging and the EMBL Genomic Core Facility for the preparation of the mRNA-Seq libraries and next generation sequencing. We are grateful to Masato Kanemaki for kindly providing us with the plasmid containing the *OsTIR1* cDNA. We also thank Javier Lizarrondo for the cartoon representation of OGT’s structure and Francesco Tabaro for invaluable support with code sharing.

## Supplementary figure legends

**Figure S1. Mutation of the putative NLS has no detectable effect on OGT nuclear localization.**

**(A)** Linear representation of the domain composition of OGT p110 (UniProt Q8CGY8), indicating in red the position of the putative nuclear localization signal DFP_461-463_ (Seo et al, 2016b). The mouse allele *Ogt^NLS-^* bears an 8 pb substitution, resulting in the DFP_461-463_->AAA_461-463_ amino acid substitution. TPR: N-terminal tetratricopeptide repeat; N/C-Catalytic: N- and C-terminal lobes of the catalytic domain.

**(B)** Representative images of OGT immunofluorescence staining (ab177941 antibody) in wild type and *Ogt^NLS-/^*^Y^ male primary MEFs, showing nuclear signal also in cells where the putative NLS is mutated. The result was confirmed using two pairs of WT and *Ogt^NLS-/^*^Y^ MEF clones. One z plane is shown. Scale bar indicates 20 μm.

**(C)** Western blot of endogenous OGT protein (ab177941 antibody) in *Ogt^NLS-/^*^Y^ and WT male primary MEFs after cellular fractionation. Vinculin (VCL) and histone H3 were probed as controls for the cytosolic and nuclear fractions, respectively. The amount of OGT^NLS-^ in the nuclear compartment is comparable to WT OGT. The result was reproduced using a different pair of WT and *Ogt^NLS-/^*^Y^ MEF clones.

**Figure S2. Deregulation of genes and retrotransposons in blastocysts with mutations of *Ogt* at T931.**

**(A)** Integrative Genomics Viewer (IGV) screenshot showing RNA-Seq reads around the T931 residue of *Ogt* for three representative genotypes of male blastocysts found in the dataset. Mutated nucleotides are colored in their containing reads or black barred when deleted. For the T931del mutant, reads marked with an asterisk were soft-clipped by STAR, which indicates that the last three mapped nucleotides (CAC) are in fact coming from the next three ones (also CAC) of the cDNA, due to the deletion. Note that amino acids 931 and 932 are both Threonines (T).

**(B)** Transcriptomes of individual male blastocysts produced by the experiment described in Fig. 2B in the space defined by PC1 and PC2 of their PCA. The variance explained by each PC is in parentheses.

**(C)** PCA of transcriptomes of individual female blastocysts produced by the experiment described in Fig. 2B.

**(D)** DESeq2-normalized counts of *Ogt* for all blastocysts produced in the experiment in Fig. 2B. P-values are from DESeq2 Wald test.

**(E)** Expression dynamics of the 10 most significant (based on DESeq2 p-value) upregulated and downregulated *Ogt^T931A/^*^Y^ DEGs throughout preimplantation development (mRNA-Seq data from GSE66582 and GSE76505). The two biological replicates per stage were averaged. For each gene, the TPM value in the E3.5 blastocyst is the highest TPM value between the values in E3.5 ICM and E3.5 trophectoderm. The mean among all genes is drawn, as well as the 95% confidence interval, computed using basic nonparametric bootstrap (R function ‘mean.cl.boot’). Y-axis ticks are in log_2_ scale. TPM: Transcript Per Million.

**(F)** PCA biplot of the FPKM expression values of the main families of retrotransposons, in male blastocysts from the experiment described in Fig. 2B. The variance explained by each PC is in parentheses and repeat families are coloured by their repeat class. N = 9 WT, 10 *Ogt^T931A/^*^Y^ embryos.

**(G)** Expression (FPKM) of the most significant (based on DESeq2 p-value) differentially expressed (adj. p-value < 0.1, any log_2_FC, mean DESeq2-normalized gene counts > 10) retrotransposons in *Ogt^T931A/^*^Y^ versus WT male blastocysts, in all blastocysts produced in the experiment described in Fig. 2B.

**(H)** Scatter plot comparing the transcriptional deregulation of retrotransposons in *Ogt^T931A/^*^Y^ versus WT male blastocysts with their expression dynamics between the 8-cell and E3.5 blastocyst stage in unperturbed embryos. The log2FC at E3.5 vs. 8-cell stage was computed as (75% log2FC E3.5 trophectoderm vs. 8-cell) + (25% log2FC E3.5 ICM vs. 8-cell). Repeats are colored by family and labeled only if significant (adj. p-value < 0.1) in the *Ogt^T931A/^*^Y^ vs. WT comparison. Pearson correlation coefficient is shown both for significantly deregulated repeats and for all repeats in the scatter plot.

**Figure S3. Developmental phenotype and deregulation of retrotransposons upon a mild reduction of OGT’s activity.**

**(A)** Average DESeq2-normalized counts of marker genes for LaTP and SynTII labyrinth clusters in single placentae with all four genotypes analyzed.

**(B)** (left) Representative images of E11.75 and E12 WT embryos among the ones used for developmental staging based on forelimb shape using the eMOSS software (https://limbstaging.embl.es/) and (right) result of the staging for all litters analyzed. N = 6 embryos dissected at E11.75 and 8 embryos dissected at E12.5 from a total of two litters from WT mothers, 16 embryos dissected at E12.5 from two litters from *Ogt^Y851A/^*^+^ mothers.

**(C)** MA-plots from DESeq2 differential expression analysis of retrotransposons in (left) *Ogt^Y851A^***-**homozygous versus heterozygous female placentae and (right) *Ogt^Y851A^***-**hemizygous versus WT male ones. All repeats with mean DESeq2-normalized gene counts > 10, adj. p-value < 0.05, any log_2_FC are colored and labeled.

**(D)** Expression (FPKM), in all four placental genotypes analyzed, of the four significantly upregulated (adj. p-value < 0.05, any log_2_FC) retrotransposons in *Ogt^Y851A/^*^Y^ versus WT male placentae together with two of the most active IAP elements (IAP-d-int and IAPEz-int, normally repressed and expressed in murine trophoblast stem cells, respectively (Weigert et al, 2023)), which are not significantly upregulated. The mean for each placental genotype is drawn. padj = adj. p-value computed using DESeq2 Wald test and corrected for multiple testing using the Benjamini and Hochberg method.

**(E)** Sum of FPKM of the only two types of satellite repeats (Repeat Library 20140131 for mm10, Dec 2011)) that are detected in the RNA-Seq dataset. They are both upregulated in male *Ogt^Y851A/^*^Y^ placentae. The mean for each placental genotype is drawn. P-value was computed using the unpaired Wilcoxon rank sum exact test on the FPKM after subtracting the sum of FPKM of all DNA repetitive elements, in order to account for DNA contamination.

In (**C**-**E**), N per genotype = 6 placentae for female homozygous and male WT, 5 for female heterozygous (due to exclusion of one outlier sample after unsupervised clustering) and male homozygous (due to exclusion of one sample with higher counts from DNA repetitive elements), all genotypes coming from at least two different litters.

**Figure S4. Rapid degradation of endogenous OGT in MEFs using the AID system.**

**(A)** Western blot detection of endogenous OGT in whole cell protein extracts from female and male E7.5 embryos WT or bearing the AID-*Ogt* allele.

**(B)** Scheme of the mouse cross used to produce double mutant E12.5 embryos (*OsTIR*,*Ogt^AID^*) and littermate controls (*OsTIR*,*Ogt^WT^*) for MEFs derivation and *in vitro* auxin-induced AID-OGT degradation.

**(C)** Quantification of the OGT western blot in Fig. 4C. The intensity of the OGT band was normalized to LAMIN C intensity. Bar plot heights and error bars show the mean and standard deviation, respectively, of two biological replicates using two pairs of *OsTIR*,*Ogt^AID^*, *OsTIR*,*Ogt^WT^* littermate clones from different litters.

**(D)** Western blot of endogenous OGT (ab177941 antibody) in whole cell protein extracts from male primary MEFs, untreated or treated with auxin for the same amount of time as for RNA-Seq analysis. Lamin A/C was probed as a loading control.

**(E)** Western blot analysis of the kinetic of O-GlcNAc depletion following OGT acute degradation in MEFs, using the RL2 anti-O-GlcNAc antibody.

**Figure S5. Inefficient O-GlcNAc perturbation using the AID-OGT degron system in the preimplantation embryo.**

**(A)** Scheme of the experiments performed to test AID-OGT degradation *ex vivo* in preimplantation embryos. Top: IVF between WT sperm and either *Ogt^WT/WT^*or *Ogt^AID/WT^ OsTIR*-expressing females was used to produce embryos untreated or treated with auxin from the fertilization plate, which were stained for OGT and O-GlcNAc at the zygote and morula stages, respectively. Bottom: natural mating of *Ogt^AID/WT^*females with *OsTIR*-homozygous males was used to produce zygotes grown *ex-vivo* until the blastocyst stage for single embryo mRNA-Seq; 24 hours before collection, half of them were moved to a medium supplemented with 250 μM auxin. In both types of experiment, females heterozygous and males homozygous AID-*Ogt* as well as control *Ogt* WT embryos, all expressing *OsTIR*, are produced for analysis.

**(B)** Test for auxin toxicity on WT embryos. Embryos were generated through IVF and cultured *ex vivo* in the absence or presence of auxin from the moment of fertilization to E4. Representative widefield microscopy images are shown for both conditions, total number of starting zygotes and percentage of E4 expanded blastocysts are indicated. Scale bar indicates 40 μm.

**(C)** DESeq2-normalized gene counts of *OsTIR1* for E4 blastocysts obtained from IVF of *OsTIR,Ogt^AID^* (N = 19 blastocysts) or control *Ogt^AID^* oocytes (not bearing the *OsTIR1* gene) (N = 15 blastocysts) with WT sperm. The mean for each group of embryos is drawn.

**(D)** Percentage of E4 blastocysts obtained from the same IVF as in (**C**). Embryos of both groups were treated with auxin from the time of fertilization to E4. Barplot heights and error bars indicate the mean and standard deviation, respectively, of three replicate experiments of IVF followed by auxin treatment. Total number of starting 2-cell embryos for the two groups is stated in the legend. P-value is for paired two-sided Student’s t-test, assuming unequal variance.

**(E)** Representative images of O-GlcNAc immunofluorescence staining (RL2 antibody) in morulae from *OsTIR*,*Ogt^AID/WT^* and control *OsTIR*,*Ogt^WT/WT^* oocytes, grown in 250 μM auxin from the time of fertilization. Embryos were mounted in drops and imaged using an X-light V3 Spinning disk confocal. One z plane is shown. Scale bar indicates 20 μm. The total number of imaged embryos is indicated.

**(F)** Quantification of the O-GlcNAc fluorescence signal from the morulae in (**A**) (Methods). Average signal per group is marked. P-value is for unpaired two-sided Student’s t-test, assuming unequal variance.

**(G)** Representative images of OGT immunofluorescence staining (ab177941 antibody) in zygotes from IVF of *OsTIR*,*Ogt^AID/WT^* oocytes with WT sperm, untreated or treated with 500 μM auxin from the time of fertilization. Embryos were mounted in slides and imaged using an X-light V3 Spinning disk confocal. One z plane is shown. Scale bar indicates 20 μm. The total number of imaged embryos is indicated.

**(H)** PCA of E4.5 blastocysts from the RNA-Seq experiment depicted in (**A**). The variance explained by each PC is in parentheses.

**(I)** For the same RNA-Seq experiment as in (**H**), MA-plots from DESeq2 differential gene expression analysis of *Ogt^AID^* versus *Ogt^WT^* genotypes’ comparison, separately for male and female blastocysts, untreated and treated with auxin. Genes with mean DESeq2-normalized gene counts > 10, adj. p-value < 0.05, any log_2_FC (DEGs) are colored, and their number is indicated. Genes with abs(log2FC) ≥ 0.2 are labeled.

In (**H**-**I**), N = 13 and 16 treated and untreated females *Ogt^AID^*, 14 and 12 treated and untreated females *Ogt^WT^*, 9 and 13 treated and untreated males *Ogt^AID^*, 11 and 11 treated and untreated males *Ogt^WT^*.

**Figure S6. Kinetics of differential gene expression after rapid degradation of endogenous OGT in MEFs.**

**(A)** Volcano plots from DESeq2 analysis of gene expression changes in (left) untreated *OsTIR*,*Ogt^AID^* versus untreated *OsTIR,Ogt^WT^* MEFs clones (effect of the hypomorphic genotype) and (right) *OsTIR,Ogt^WT^* control clones treated with auxin for 96 hours versus grown untreated (effect of the auxin drug). Differentially expressed genes with p-value < 10^-5^ and absolute log_2_FC > 0.3 (i.e. 1.2-fold increase or decrease in expression) are labeled in red and their number is indicated.

**(B)** DESeq2-normalized counts of *Ogt*, *Oga* and the transgene *OsTIR1*. *Oga* level was already downregulated in the untreated *OsTIR*,*Ogt^AID^* genotype, which is *de facto* an hypomorphic *Ogt* mutant, when compared with the *OsTIR*,*Ogt^WT^* one. Note that *OsTIR1* is also significantly downregulated after auxin addition, suggesting that an unknown mechanism counteracts the degron system upon auxin treatment. Mean of counts for each group of samples is shown. Y-axis ticks are in log_10_ scale. padj = adj. p-value computed using DESeq2 Wald test and corrected for multiple testing using the Benjamini and Hochberg method.

**(C)** Number and overlap of differentially expressed genes (DEGs; adj. p-value < 0.05, any log_2_FC) in auxin-treated *OsTIR,Ogt^AID^*clones versus untreated *OsTIR,Ogt^AID^* clones at the three time points analyzed. DEGs in auxin-treated control clones at any time point are excluded.

**(D)** Gene ontology (GO) over-representation analysis of DEGs (adj. p-value < 0.05, any log_2_FC) for the same comparisons between genotypes and conditions as in Fig. 4E, but showing result for Cellular Component (CC) and Molecular Function (MF) GO terms. The first 25 most enriched CC and MF GO terms are shown, based on p-value across all comparisons. Terms are ordered by gene ratio. Gene ratio = genes belonging to the GO term / total number of deregulated DEGs for that comparison. UP = upregulated DEGs, DOWN = downregulated DEGs. Terms enriched due to auxin treatment on control clones are written in gray.

