## Supplemental Figures for "Genetic gradual reduction of OGT activity unveils the essential role of O-GlcNAc in the mouse embryo"

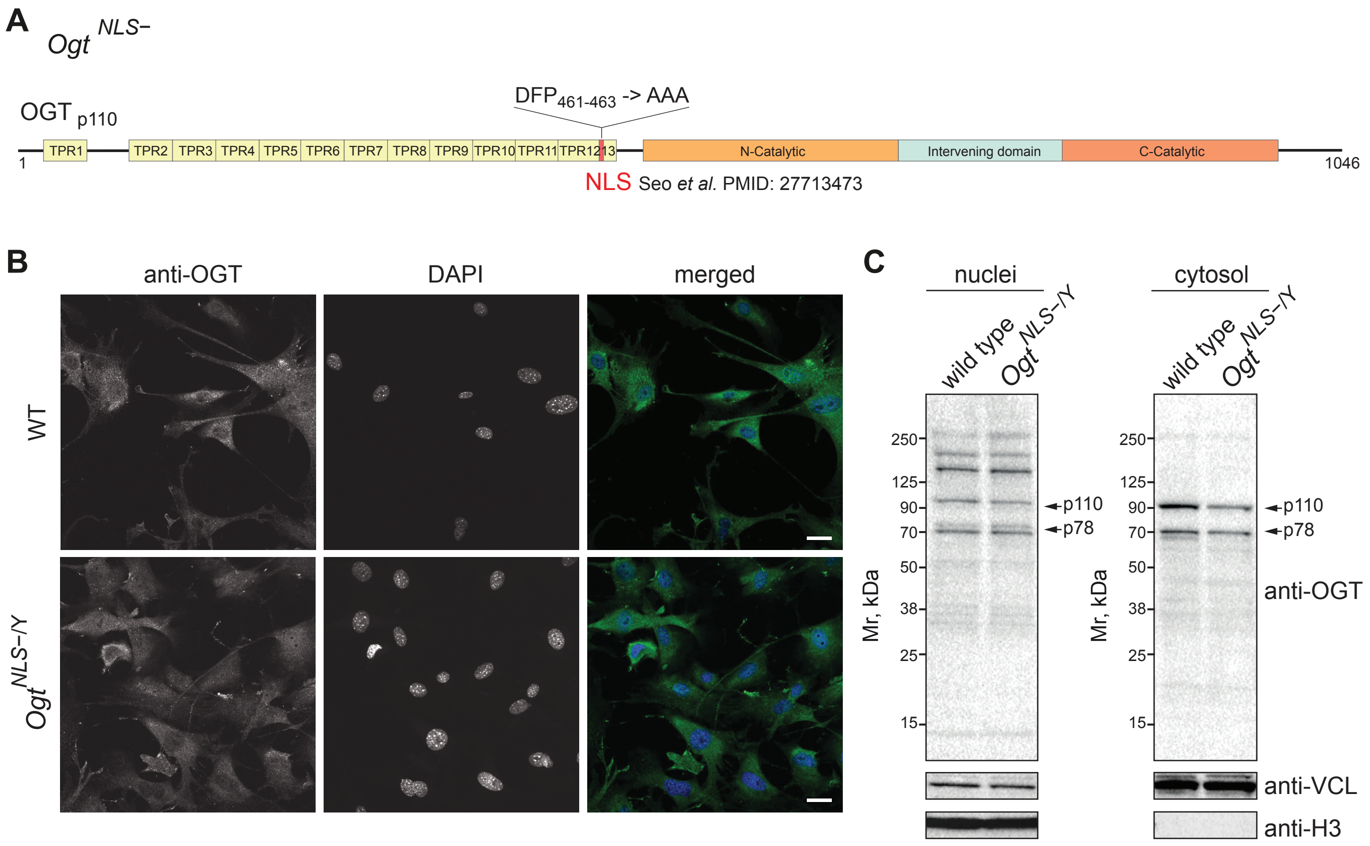


**Figure S1** Mutation of the putative NLS has no detectable effect on OGT nuclear localization.

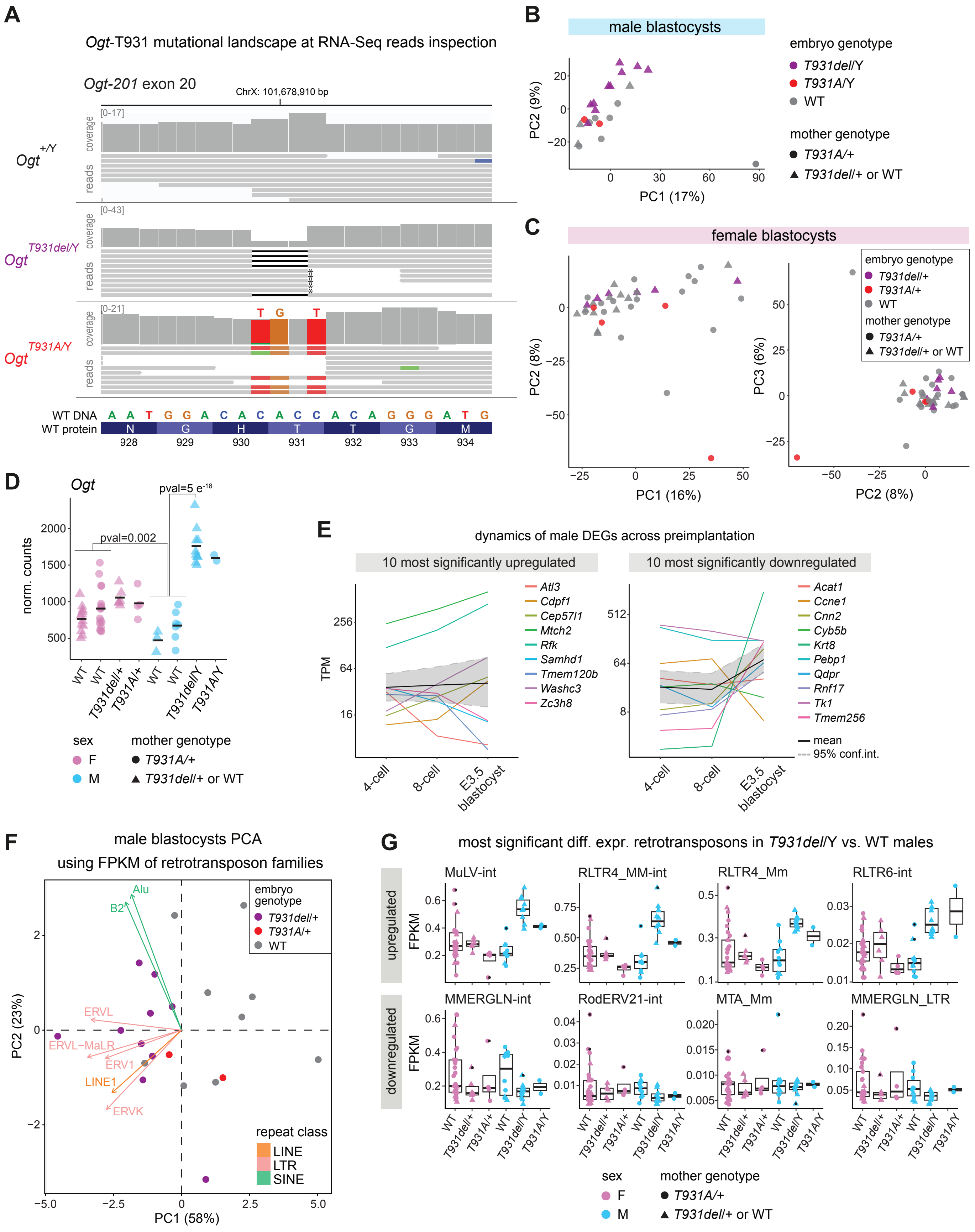


**Figure S2** Deregulation of genes and retrotransposons in blastocysts with mutations of *Ogt* at T931.

H

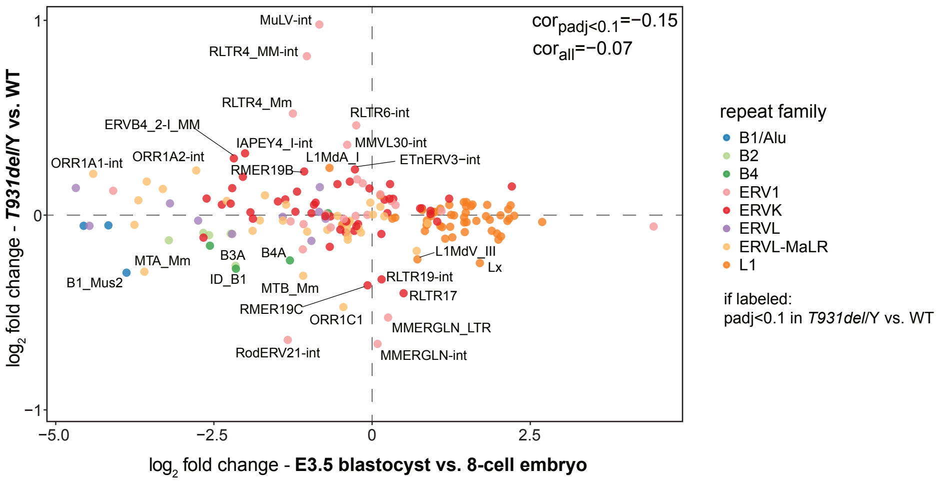


**Figure S2** *(continued)*.

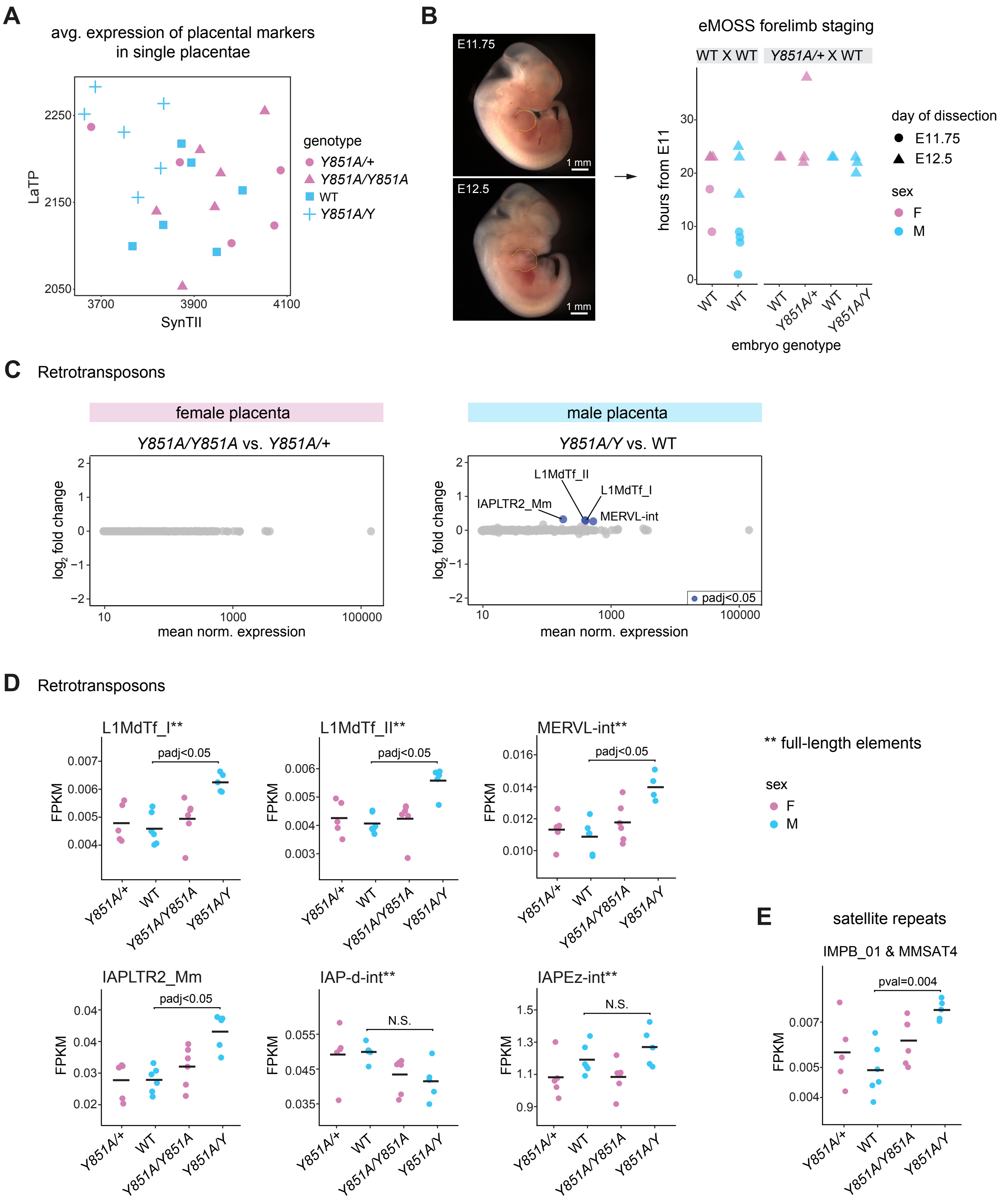


**Figure S3** Developmental phenotype and deregulation of retrotransposons upon a mild reduction of OGT's activity.

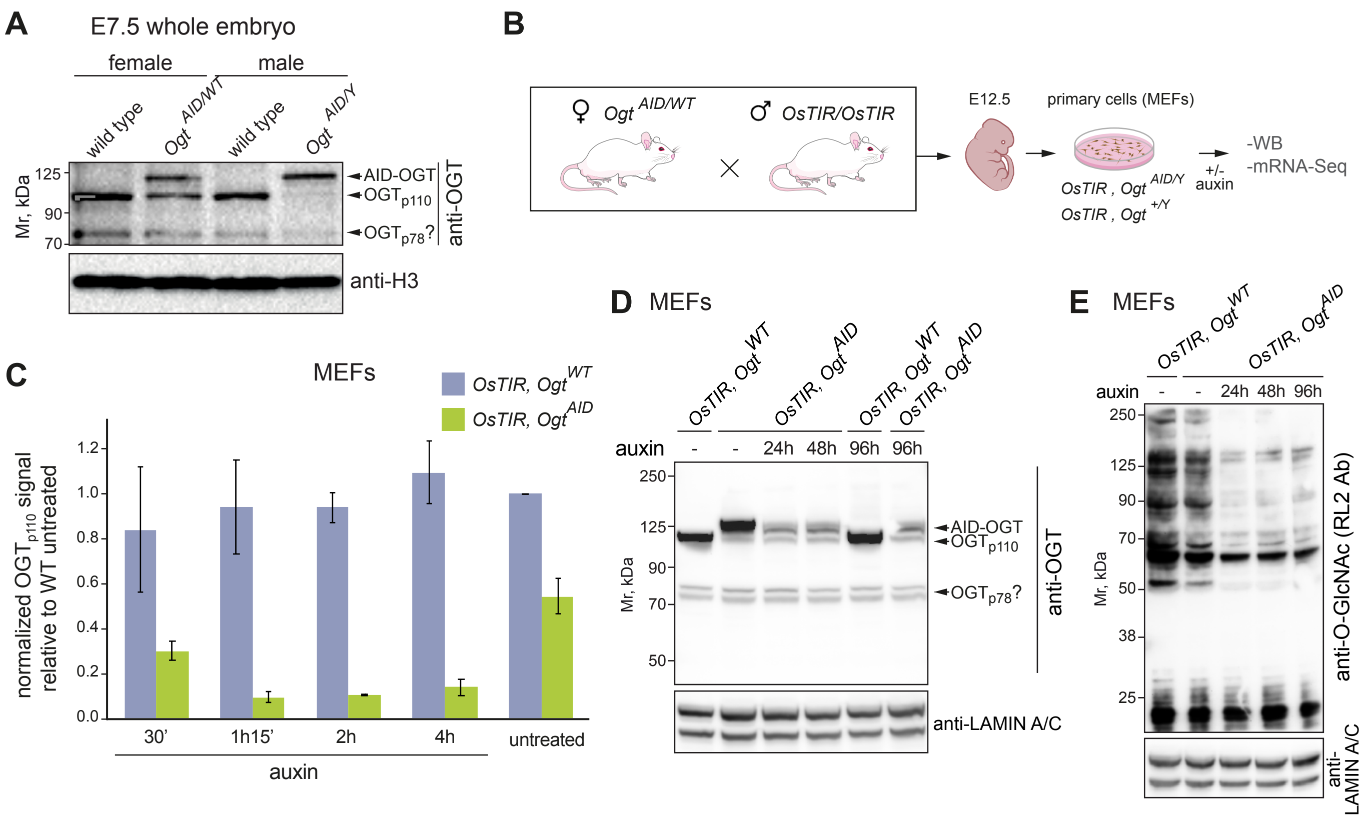


**Figure S4** Rapid degradation of endogenous OGT in MEFs using the AID system.

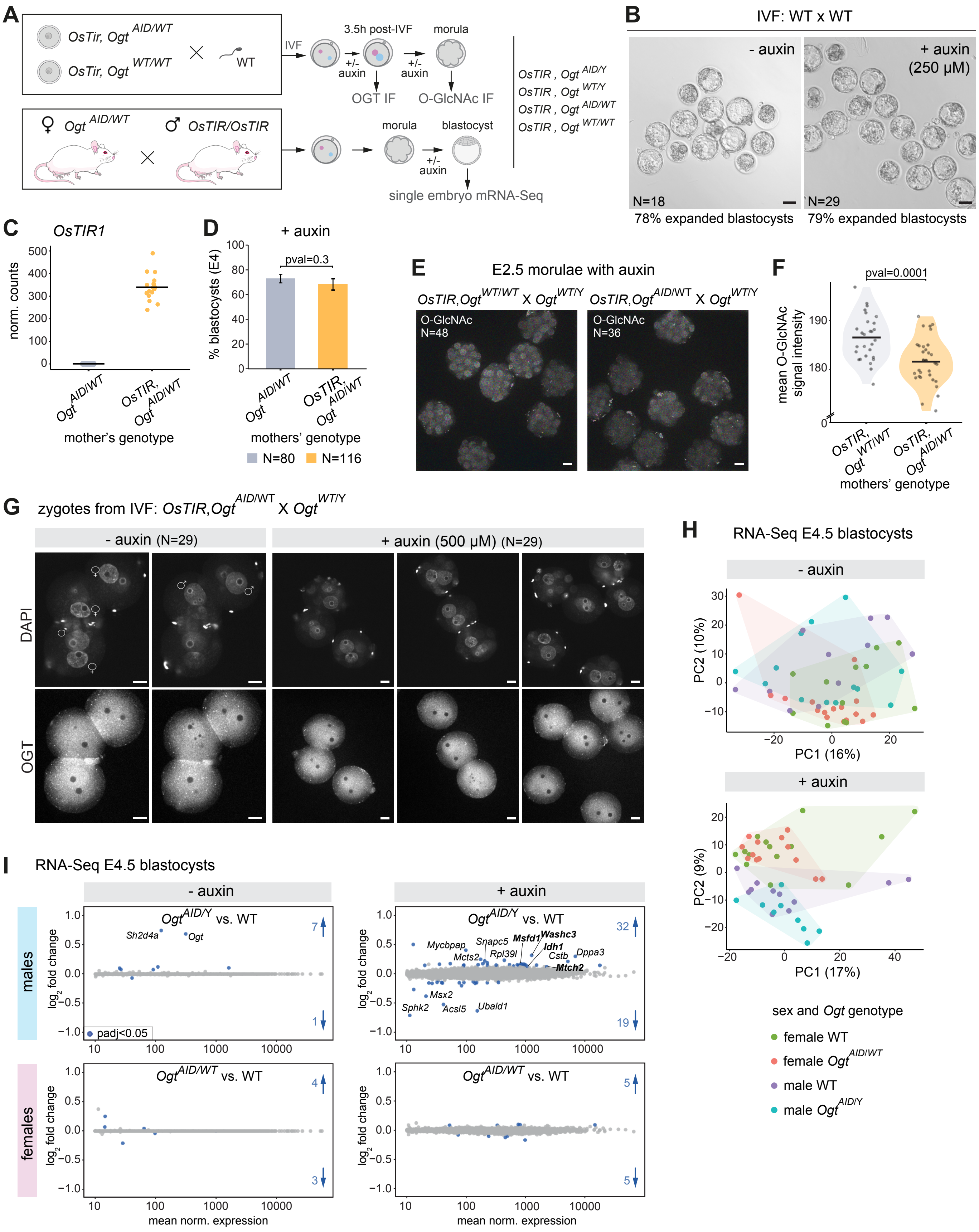


**Figure S5** Inefficient O-GlcNAc perturbation using the AID-OGT degron system in the preimplantation embryo.

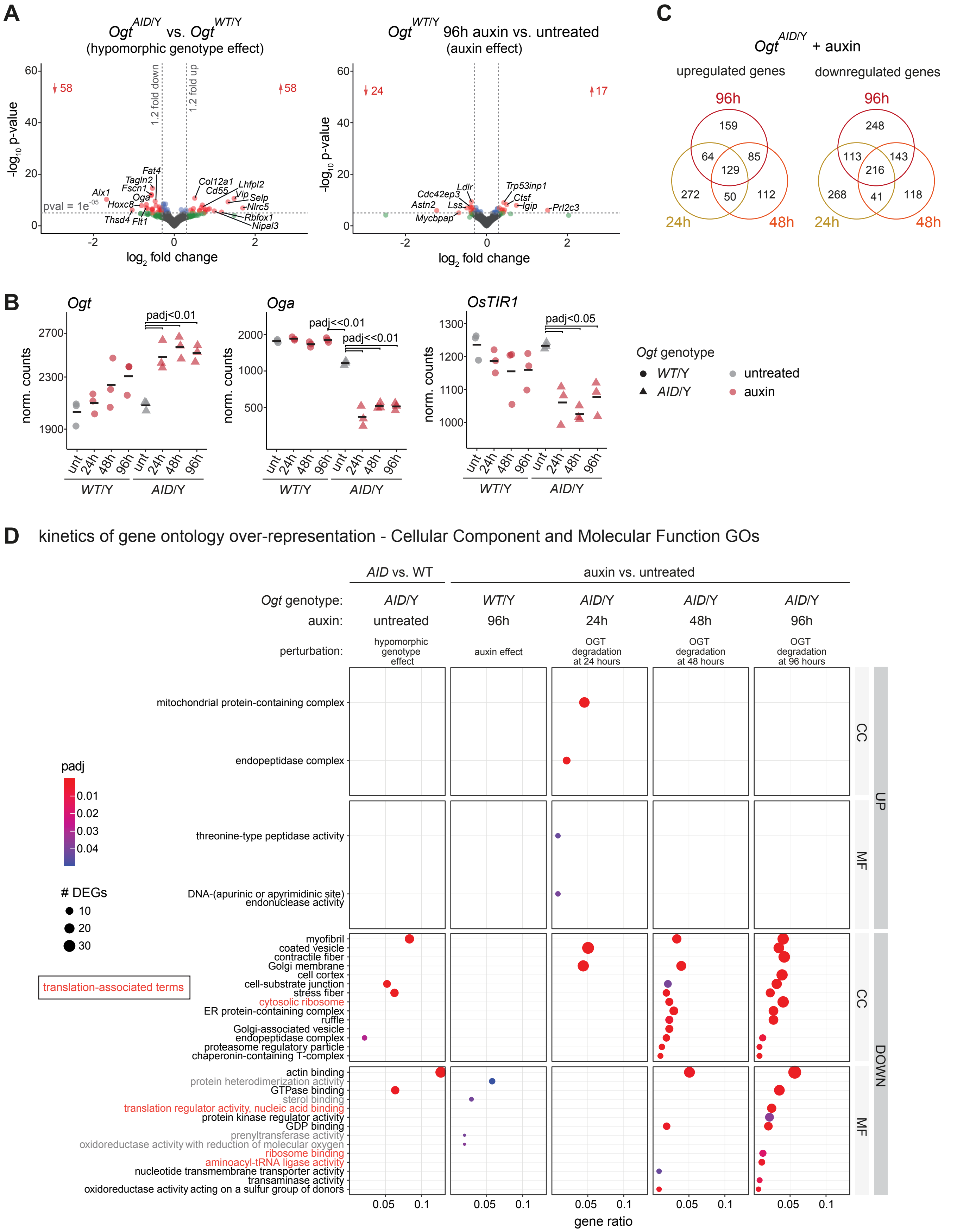


**Figure S6** Kinetics of differential gene expression after rapid degradation of endogenous OGT in MEFs.
