## Supplemental Table 1 for "Genetic gradual reduction of OGT activity unveils the essential role of O-GlcNAc in the mouse embryo"

Table S1: list of generated mouse alleles

| **Allele name** | | **Mutation type** | **Targeted exon** | **Genomic coordinates insertion or substitution (GRCm39/mm39)** |
|---|---|---|---|---|
| Short name | Standardized nomenclature | | | |
| *Ogt*$^{Y851A}$ | FVB/NCrl-*Ogt*$^{em1(Y851A)}$Emr | substitution Y851A | exon 19 (*Ogt-201*) | substitution: GTACTGTAACTTTAATCAGTTATATAAAATTGACCCATCT -> CTATTGCAATTTCAACCAACTGGCCAAGATCGATCCTAGC, ChrX:100719847->100719886 forward strand |
| *Ogt*$^{T931A}$ | FVB/NCrl-*Ogt*$^{em2(T931A)}$Emr | substitution T931A | exon 20 (*Ogt-201*) | substitution: CACC -> TGCT, ChrX:100722515->100722518 forward strand |
| *Ogt*$^{Q949N}$ | FVB/NCrl-*Ogt*$^{em3(Q949N)}$Emr | substitution Q949N | exon 19 (*Ogt-201*) | substitution: GTACTGTAACTTTAATCAGTTATATAAAATTGACCCATCT -> CTATTGCAATTTCAACAACCTGTACAAGATCGATCCTAGC, ChrX:100719847->100719886 forward strand |
| *Ogt*$^{H568A}$ | FVB/NCrl-*Ogt*$^{em4(H568A)}$Emr | substitution H568A | exon 13 (*Ogt-201*) | substitution: GAATCAC -> GAATGCT, ChrX:100713458->100713464 forward strand |
| *Ogt*$^{NLS-}$ | FVB/NCrl-*Ogt*$^{em5(DFPmut)}$Emr | substitution DFP(461-463) -> AAA | exon 11 (*Ogt-201*) | substitution: GACTTTCCT -> GCTGCTGCA, ChrX:100711182->100711190 forward strand |
| *Ogt*$^{NterAID-MYC-FLAG}$ | FVB/NCrl-*Ogt*$^{em6(AID)}$Emr | insertion [N-ter AID-MYC-FLAG] | exon 1 (*Ogt-201*) | insertion: ChrX:100683892 forward strand CCTAAAGACCCTGCAAAACCTCCCGCAAAGGCACAGGTGGTTGGATGGCCCCCAGTGAGATCCTATCGAAAGAACGTAATGGTTTCTTGTCAAAAGAGTAGTGGTGGACCTGAGGCCGCAGCCTTCGTGAAGGAACAAAAACTTATCAGCGAGGAAGACCTCGAGCAGAAGCTGATCAGCGAGGAAGACCTGGATTATAAAGACGACGATGATAAA |

| | | | | |
|---|---|---|---|---|
| ROSA26$^{OsTIR}$ | FVB;B6J;129-Gt(ROSA)26Sor$^{tm1(OsTIR)}$Emr | Insertion [OsTIR-Myc-HA] | ROSA26 locus | Insertion: Chr6:113,076,033 (mm10):<br>ATCTGTAGGGCGCAGTAGTCCAGGG<br>TTTCCTTGATGATGTCATACTTATCCT<br>GTCCCTTTTTTTTCCACAGCTCGCGG<br>TTGAGGACAAACTCTTCGCGGTCTT<br>TGTGCACTTAAGATAACTTCGTATAG<br>CATACATTATACGAAGTTATCCAGTG<br>GGGATCGACGGTATCGATAAGCTTCC<br>ACCATGACATACTTTCCTGAAGAGGT<br>CGTCGAACACATTTTTAGCTTCCTGC<br>CTGCACAGAGAGATAGAAACACAGT<br>GAGCCTGGTCTGCAAAGTGTGGTAC<br>GAGATCGAACGCCTGAGCCGGAGAG<br>GAGTGTTCGTCGGCAACTGCTATGCT<br>GTGAGAGCAGGCAGGGTCGCCGCTA<br>GGTTTCCAAATGTGCGCGCACTGAC<br>CGTCAAGGGGAAACCCCACTTCGCC<br>GACTTTAACCTGGTGCCCCCTGATTG<br>GGGAGGATACGCCGGCCCTTGGATC<br>GAGGCAGCCGCTCGCGGCTGTCATG<br>GACTGGAGGAACTGCGCATGAAGCG<br>AATGGTGGTCTCTGACGAAAGTCTG<br>GAGCTGCTGGCTCGGAGCTTCCCTA<br>GGTTTCGCGCACTGGTGCTGATTTCT<br>TGCGAAGGCTTCAGCACCGATGGAC<br>TGGCAGCCGTGGCCTCCCACTGTAA<br>GCTGCTGCGGGAGCTGGACCTCCAG<br>GAGAATGAAGTGGAGGATAGAGGCC<br>CCAGATGGCTGTCTTGCTTCCCAGAC<br>TCATGTACCAGCCTGGTGTCCCTGAA<br>CTTTGCCTGCATCAAAGGCGAAGTG<br>AATGCTGGGTCCCTGGAGCGGCTGG<br>TCTCAAGAAGCCCCAACCTGAGGTC<br>TCTGCGGCTGAACCGGAGCGTGAGC<br>GTGGACACTCTGGCTAAGATTCTGCT<br>GAGAACCCCTAACCTGGAGGATCTG<br>GGAACCGGCAATCTGACAGACGATT<br>TCCAGACAGAATCCTACTTTAAACTG<br>ACTTCTGCCCTGGAGAAGTGTAAAA<br>TGCTGAGGAGTCTGTCAGGATTCTG<br>GGATGCTTCACCCGTGTGCCTGAGC<br>TTTATCTACCCTCTGTGTGCACAGCT<br>GACAGGCCTGAACCTGAGCTATGCA |

| | | | | |
|---|---|---|---|---|
| | | | | CCAACCCTGGACGCCAGTGATCTGA CAAAGATGATCTCACGCTGCGTGAA ACTCCAGCGACTGTGGGTGCTGGAC TGTATTTCCGATAAGGGGCTCCAGGT GGTCGCCAGCTCCTGCAAGGACCTC CAGGAGCTGAGAGTGTTCCCATCTG ATTTTTACGTGGCCGGATATAGTGCT GTCACTGAGGAAGGCCTGGTGGCAG TCTCACTGGGATGCCCAAAGCTGAA CAGCCTGCTGTATTTCTGTCATCAGA TGACTAATGCTGCACTGGTGACCGTC GCCAAGAACTGCCCTAATTTCACCC GATTTCGGCTGTGTATTCTGGAACCA GGCAAACCCGACGTGGTCACATCCC AGCCACTGGATGAAGGGTTTGGAGC TATCGTGAGAGAGTGCAAGGGACTC CAGAGGCTGAGCATTTCCGGCCTGC TGACAGACAAAGTGTTCATGTACATC GGCAAGTATGCTAAGCAGCTGGAGA TGCTGAGCATTGCATTTGCCGGAGA CTCCGATAAGGGCATGATGCACGTG ATGAACGGGTGTAAGAATCTGCGAA AACTGGAAATCCGGGACAGCCCTTT CGGGGATGCCGCTCTGCTGGGAAAC TTTGCCAGATACGAGACAATGAGGA GCCTGTGGATGTCTAGTTGCAATGTG ACTCTGAAGGGCTGTCAGGTCCTGG CTAGTAAAATGCCTATGCTGAACGTG GAAGTCATTAATGAGCGGGACGGGT CTAACGAAATGGAGGAAAATCATGG CGACCTGCCAAAGGTGGAGAAACTG TATGTGTATCGGACCACCGCAGGGG CAAGAGATGATGCTCCCAACTTTGT GAAGATTCTGGAGGAGCAGAAGCTG ATCTCAGAGGAGGACCTGTACCCAT ACGATGTTCCAGATTACGCTGTCGAC TAGCATATGTACGAAGTTATAAGCTG GAAGTTCCTATTCTCTAGAAAGTATA GGAACTTCAAGCTTAGGTGGCACTT TTCGGGCTACCGGGTAGGGGAGGCG CTTTTCCCAAGGCAGTCTGGAGCAT GCGCTTTAGCAGCCCCGCTGGGCAC TTGGCGCTACACAAGTGGCCTCTGG |

| | | | | |
|---|---|---|---|---|
| | | | | CCTCGCACACATTCCACATCCACCGG<br>TAGGCGCCAACCGGCTCCGTTCTTTG<br>GTGGCCCCTTCGCGCCACCTTCTACT<br>CCTCCCCTAGTCAGGAAGTTCCCCCC<br>CGCCCCGCAGCTCGCGTCGTGCAGG<br>ACGTGACAAATGGAAGTAGCACGTC<br>TCACTAGTCTCGTGCAGATGGACAG<br>CACCGCTGAGCAATGGAAGCGGGTA<br>GGCCTTTGGGGCAGCGGCCAATAGC<br>AGCTTTGCTCCTTCGCTTTCTGGGCT<br>CAGAGGCTGGGAAGGGGTGGGTCC<br>GGGGGCGGGCTCAGGGGCGGGCTC<br>AGGGGCGGGGCGGGCGCCCGAAGG<br>TCCTCCGGAGGCCCGGCATTCTGCA<br>CGCTTCAAAAGCGCACGTCTGCCGC<br>GCTGTTCTCCTCTTCCTCATCTCCGG<br>GCCTTTCGACCTGCAGCAGCACGTG<br>TTGACAATTAATCATCGGCATAGTATA<br>TCGGCATAGTATAATACGACAAGGTG<br>AGGAACTAAACCATGGGATCGGCCA<br>TTGAACAAGATGGATTGCACGCAGG<br>TTCTCCGGCCGCTTGGGTGGAGAGG<br>CTATTCGGCTATGACTGGGCACAACA<br>GACAATCGGCTGCTCTGATGCCGCC<br>GTGTTCCGGCTGTCAGCGCAGGGGC<br>GCCCGGTTCTTTTTGTCAAGACCGA<br>CCTGTCCGGTGCCCTGAATGAACTG<br>CAGGACGAGGCAGCGCGGCTATCGT<br>GGCTGGCCACGACGGGCGTTCCTTG<br>CGCAGCTGTGCTCGACGTTGTCACT<br>GAAGCGGGAAGGGACTGGCTGCTAT<br>TGGGCGAAGTGCCGGGGCAGGATCT<br>CCTGTCATCTCACCTTGCTCCTGCCG<br>AGAAAGTATCCATCATGGCTGATGCA<br>ATGCGGCGGCTGCATACGCTTGATCC<br>GGCTACCTGCCCATTCGACCACCAA<br>GCGAAACATCGCATCGAGCGAGCAC<br>GTACTCGGATGGAAGCCGGTCTTGT<br>CGATCAGGATGATCTGGACGAAGAG<br>CATCAGGGGCTCGCGCCAGCCGAAC<br>TGTTCGCCAGGCTCAAGGCGAGCAT<br>GCCCGACGGCGAGGATCTCGTCGTG<br>ACCCATGGCGATGCCTGCTTGCCGA |

| | | | | |
|---|---|---|---|---|
| | | | | ATATCATGGTGGAAAATGGCCGCTTT<br>TCTGGATTCATCGACTGTGGCCGGCT<br>GGGTGTGGCGGACCGCTATCAGGAC<br>ATAGCGTTGGCTACCCGTGATATTGC<br>TGAAGAGCTTGGCGGCGAATGGGCT<br>GACCGCTTCCTCGTGCTTTACGGTAT<br>CGCCGCTCCCGATTCGCAGCGCATC<br>GCCTTCTATCGCCTTCTTGACGAGTT<br>CTTCTGAGCGGGACTCTGGGGTTCG<br>AAATGACCGACCAAGCGACGCCCAA<br>CCTGCCATCACGAGATTTCGATTCCA<br>CCGCCGCCTTCTATGAAAGGTTGGG<br>CTTCGGAATCGTTTTCCGGGACGCC<br>GGCTGGATGATCCTCCAGCGCGGGG<br>ATCTCATGCTGGAGTTCTTCGCCCAC<br>CCTAGGGGGAGGCTAACTGAAACAC<br>GGAAGGAGACAATACCGGAAGGAA<br>CCCGCGCTATGACGGCAATAAAAAG<br>ACAGAATAAAACGCACGGTGTTGGG<br>TCGTTTGTTCATAAACGCGGGGTTCG<br>GTCCCAGGGCTGGCACTCTGTCGAT<br>ACCCCACCGAGACCCCATTGGGGCC<br>AATACGCCCGCGTTTCTTCCTTTTCC<br>CCACCCCACCCCCCAAGTTCGGGTG<br>AAGGCCCAGGGCTCGCAGCCAACGT<br>CGGGGCGGCAGGCCCTGCCAGGATC<br>CGAAGTTCCTATTCTCTAGAAAGTAT<br>AGGAACTTCCTCGAGTTTAAACTAA<br>GCTGATCAGCCTCGACTGTGCCTTCT<br>AGTTGCCAGCCATCTGTTGTTTGCCC<br>CTCCCCCGTGCCTTCCTTGACCCTGG<br>AAGGTGCCACTCCCACTGTCCTTTCC<br>TAATAAAATGAGGAAATTGCATCGCA<br>TTGTCTGAGTAGGTGTCATTCTATTC<br>TGGGGGGTGGGGTGGGGCAGGACA<br>GCAAGGGGGAGGATTGGGAAGACA<br>ATAGCAGGCATGCTGGGGATGCGGT<br>GGGCTTATGGCTTCTGAGGCGGAAA<br>GAACCAGCTGGGGCTCGAT |
