## Supplemental Table 2 for "Genetic gradual reduction of OGT activity unveils the essential role of O-GlcNAc in the mouse embryo"

Table S2: primers for PCR genotyping of mouse lines

| Gene target | Primer direction | Sequence (5'-3') |
|---|---|---|
| *Ogt*$^{NterAID\text{-}MYC\text{-}FLAG}$ | forward | AGTAGTGGCGGCAGTAGAAG |
| | reverse | TAATGGGGATGGTCAGAGGG |
| *OsTIR* insert (to genotype for the presence of the insert at least on one allele) | forward | AGAGATAGAAACACAGTGAGCC |
| | reverse | TCGCAAGAAATCAGCACCAG |
| *OsTIR* flanking sequence (to genotype for homozygosity, if used together with primers in row above) | forward | AGTCGCTCTGAGTTGTTATCAG |
| | reverse | AGGTTAGCCTTTAAGCCTGC |
| *Ogt*$^{Q949N}$ | forward | CTTGACTCAAAACCAGGGCC |
| *Ogt*$^{Q949N}$ | reverse | ATGGGGAAGGGAGATTCAGC |
| *Ogt*$^{Y851A}$ | forward | CTTGACTCAAAACCAGGGCC |
| | reverse | ATGGGGAAGGGAGATTCAGC |
| *Ogt*$^{+}$ allele and *Ogt*$^{T931A}$ allele at the region containing T931* | reverse | CATGTGGTCAGGTTTGTTGC |
| | forward | GCGTTTTCCAGCAGTAGGA |
| only *Ogt*$^{T931A}$ allele* | reverse | GAACATCCATCCCTGTAGCA |
| | forward | GCGTTTTCCAGCAGTAGGA |

* both pairs of primers were used for genotyping of the *Ogt*$^{T931A}$ mouse line: a positive signal with the first pair is necessary to interpret a negative signal with the second pair as a true wild type.
