## Supplemental Table 3 for "Genetic gradual reduction of OGT activity unveils the essential role of O-GlcNAc in the mouse embryo"

Table S3: primers for PCR genotyping of MEFs

| **Gene target** | **Primer direction** | **Sequence (5'-3')** |
|---|---|---|
| *Ogt*$^{NterAID\text{-}MYC\text{-}FLAG}$ | forward | AGTAGTGGCGGCAGTAGAAG |
| | reverse | TAATGGGGATGGTCAGAGGG |
| *OsTIR* | forward | AGAGATAGAAACACAGTGAGCC |
| | reverse | TCGCAAGAAATCAGCACCAG |
| *Sly-Xlr* (McFarlane et al., 2013) | forward | GATGATTTGAGTGGAAATGTGAGGTA |
| | reverse | CTTATGTTTATAGGCATGCACCATGTA |
