## Supplemental Table 4 for "Genetic gradual reduction of OGT activity unveils the essential role of O-GlcNAc in the mouse embryo"

Table S4: primers used for PCR genotyping of *Ogt*$^{T931A}$ blastocysts

| Gene target | Primer direction | Sequence (5'-3') |
|---|---|---|
| Xist (cDNA) | forward | TCTATCTTGTGGGTCCTGGAG |
| | reverse | CTCCTCTAAATCCAGGCAATCC |
| Ddx3y (cDNA) | forward | TGGAGGAGGAAATACAGAGAGC |
| | reverse | GGAGGACAATTATTTCCAGTTGC |
| Eif2s3y (cDNA) | forward | TGGCTGTGAAGTTGATGACC |
| | reverse | CCTTCTGTACGTACACCTAGG |
| *Ogt*$^{+}$ allele and *Ogt*$^{T931A}$ allele* | reverse | CATGTGGTCAGGTTTGTTGC |
| | forward | GCGTTTTCCAGCAGTAGGA |
| only *Ogt*$^{T931A}$ allele* | reverse | GAACATCCATCCCTGTAGCA |
| | forward | GCGTTTTCCAGCAGTAGGA |

* both pairs of primers were used for genotyping of the *Ogt*$^{T931A}$ blastocysts: a positive signal with the first pair is necessary to interpret a negative signal with the second pair as a true wild type.
