## Supplemental Table 5 for "Genetic gradual reduction of OGT activity unveils the essential role of O-GlcNAc in the mouse embryo"

Table S5: details on the generation and filtering steps of the single embryo Smart-Seq datasets

| **Mouse line** | **Type of sequencing** | **# pooled embryos in sequencing run** | **Average # of reads per embryo** | **# pooled embryos in sequencing run, per condition** | | **# embryos discarded for < $10^6$ reads** | **# embryos discarded for high rDNA /mt gene counts** | **# embryos discarded for uncertain genotype** | **# embryos discarded for visible batch effect** | **# embryos for DE analysis** |
|---|---|---|---|---|---|---|---|---|---|---|
| *Ogt*$^{NterAID\text{-}MYC\text{-}FLAG}$ | 75 bp SE | 119 | $4.5*10^6$ | AUX | 60 | 2 | 0 | 1 | 10 | 47 |
| | | | | UNT | 59 | 0 | 0 | 0 | 7 | 52 |
| *Ogt*$^{T931A}$ | 40 bp PE | 71 | $8.6*10^6$ | WT mothers | 37 | 5 | 0 | 1 | Not applicable | 31 |
| | | | | *Ogt*$^{T931A/+}$ mothers | 34 | 4 | 0 | 0 | Not applicable | 30 |
